# Interrogating the plasma proteome of repetitive head impact exposure and chronic traumatic encephalopathy

**DOI:** 10.1101/2024.10.01.616097

**Authors:** Rowan Saloner, Kaitlin B. Casaletto, Sruti Rayaprolu, Paramita Chakrabarty, Jose F. Abisambra, Salvatore Spina, Lea T. Grinberg, William W. Seeley, Bruce L. Miller, Joel H. Kramer, Gil D. Rabinovici, Breton M. Asken

## Abstract

**Background:** Exposure to repetitive head impacts (RHI) is associated with increased risk for chronic traumatic encephalopathy (CTE), a neurodegenerative tauopathy, and other neuropathological changes. Biological drivers of RHI-related neurodegeneration are not well understood. We interrogated the plasma proteome in aging adults with prior RHI compared to healthy controls (CTL) and individuals with Alzheimer’s disease (AD), including a subset characterized neuropathologically at autopsy.

**Methods:** Proximity extension assay (Olink Explore®) quantified 2,779 plasma proteins in 22 RHI patients (all AD-biomarker negative), 39 biomarker-confirmed AD, and 44 CTL. A subset of participants went to autopsy (N=16) allowing for comparisons of the antemortem plasma proteome between autopsy-confirmed CTE+ (N=7) and CTE-(N=9). Differential abundance and co-expression network analyses identified plasma proteomic signatures of RHI, which were functionally annotated using gene ontology and cell type enrichment analysis. Nonparametric correlations examined plasma proteomic associations with orthogonally-measured plasma biomarkers, global cognitive function, and semi-quantitative ratings of neuropathology burden at autopsy.

**Results:** Differential abundance analysis revealed 434 increased (vs. 6 decreased) proteins in RHI vs. CTL and 193 increased (vs. 14 decreased) in RHI vs. AD. Network analysis identified 9 protein co-expression modules (M1-M9), of which 7 were elevated in RHI compared to AD or CTL. Modules with increased abundance in RHI were enriched for mitochondrial/metabolic, cell division, and immunovascular (e.g., cell adhesion, TNF-signaling) processes. RHI-related modules exhibited strong and selective correlations with immunoassay-based plasma IL-6 in RHI cases, including the M2 TNF-signaling/cell adhesion module which harbored proteins that strongly tracked with cognitive function. RHI-related plasma protein signatures were similar in the subset of participants with autopsy-confirmed CTE, including immune and metabolic modules that positively correlated with medial temporal lobe tau and TDP-43 burden.

**Conclusions:** Molecular pathways in plasma most consistently implicated in RHI were tied to immune response, mitochondrial function, and cell metabolism. RHI-related proteomic signatures tracked with antemortem cognitive severity and postmortem neuropathological burden, providing converging evidence for their role in disease progression. Differentially abundant proteins and co-expression modules in RHI may inform mechanisms linking RHI to increased dementia risk, thus guiding diagnostic biomarker and therapeutic development for at-risk populations.

## Background

Head trauma, including traumatic brain injuries and repetitive head impacts (RHI), is an established dementia risk factor(1). RHI is a unique environmental exposure usually related to contact and collision sport participation that predisposes to multiple and mixed proteinopathies(2, 3). Chronic traumatic encephalopathy (CTE) is a progressive neurodegenerative tauopathy highly specific to prior RHI(4–6), but several other neurodegenerative diseases are linked to RHI. Lewy body diseases (e.g., Parkinson disease) and TDP43 proteinopathies (e.g., amyotrophic lateral sclerosis, frontotemporal dementias) are more common in individuals with prior RHI, and up to 80% of brain donors with CTE have at least one of these co-pathologies(7–12). Data relating RHI to increased risk for developing AD pathology are less compelling(13–15). Understanding the biological drivers of RHI-related neurodegeneration has important implications for dementia prevention, risk modification, and intervention for millions of at-risk older adults with prior RHI from popular activities like contact-collision sport play and from other sources like military service or intimate partner violence(16).

Studying neurodegenerative consequences of RHI in living adults is challenging because of highly variable symptoms and potential underlying etiologies. Research criteria for traumatic encephalopathy syndrome (TES) aim to aid in identifying living individuals with CTE based on extent of RHI exposure and symptom profiles, anchored to the common complaints of memory loss, executive dysfunction, and neurobehavioral dysregulation (i.e., explosivity, short fuse) among brain donors with CTE(17). Brain imaging studies support susceptibility of medial temporal and limbic network structures to the effects of RHI(3, 18–21), but with diverse neuropathological changes observed in these regions at autopsy (e.g., CTE, TDP-43, white matter disease)(3, 20, 22, 23). The molecular underpinnings of neuropathological change and clinical sequelae among people with RHI are less understood. Identifying biological signatures of RHI in-vivo could inform diagnostic biomarker and therapeutic development for CTE and shed light on shared mechanisms underlying increased risk for multiple neurodegenerative diseases.

Large-scale proteomics in postmortem cortical tissue of CTE brain donors has identified increased abundance of protein communities linked to inflammation, glial cell proliferation, immunoglobulins, and extracellular matrix integrity(24). Other brain tissue proteomics work studying multiple neurodegenerative tauopathies has revealed both shared and unique disease-associated microglial, endothelial, and mitochondrial protein changes across tauopathies, including CTE(25). While limited by reduced central nervous system specificity, plasma proteomics has identified brain-relevant molecular changes that are detectable in blood of living patients with neurodegenerative diseases(26). Plasma proteomics importantly increases scalability and representation of participants collected, supporting robust replication and validation of hypothesis testing in-vivo. Although targeted plasma biomarker analysis implicates inflammatory changes in RHI (e.g., elevated plasma IL-6) compared to both healthy controls and individuals with Alzheimer’s disease (AD)(19), unbiased plasma proteomics has not been applied to clinically symptomatic, living individuals with prior RHI.

We employed a high-throughput proximity-extension assay (Olink) to interrogate the plasma proteome in a well characterized clinical cohort with substantial prior RHI, most of whom met research criteria for TES. To inform proteomic changes unique to RHI, we compared RHI cases to a clinically normal sample of older adults without RHI (AD biomarker negative) and a cognitively impaired group without RHI and biomarker-confirmed AD. A subset of participants underwent brain autopsy, allowing us to compare brain donors with and without neuropathologically-confirmed CTE. Using differential abundance and weighted gene correlation network analyses (WGCNA), we identified large-scale plasma protein communities and individual biomarker candidates uniquely dysregulated in RHI, mirrored in a subset with autopsy-confirmed CTE, and correlated with orthogonally-measured plasma biomarkers, cognitive function, and neuropathological burden of tau and TDP43.

## Methods

### UCSF Memory and Aging Center

Plasma samples were collected from participants enrolled in research studies at the University of California, San Francisco (UCSF) Memory and Aging Center, either the UCSF Alzheimer’s Disease Research Center (ADRC) or cognitively unimpaired controls from the Brain Aging Network for Cognitive Health (BrANCH), as previously described(19). All participants underwent clinical evaluations including comprehensive history, neurologic exam, neuropsychological testing, caregiver interview and functional assessment (Clinical Dementia Rating scale; CDR), and blood draw. Consensus diagnoses were provided by a multidisciplinary team.

RHI participants had known RHI exposure through contact or collision sport participation or, if available, a neuropathological diagnosis of CTE. Among RHI, TES criteria and levels of diagnostic certainty were retrospectively applied using the updated 2021 research framework(17). As most, but not all, RHI participants met TES criteria, we refer to this group as “RHI” throughout. RHI participants in our cohort were all male and predominantly included former American football players. For primary clinical grouping, all RHI participants had a negative Aβ-PET scan. Two RHI participants with positive Aβ-PET scans but autopsy-confirmed CTE were excluded from primary analyses but were included in CTE-focused sub analyses (see neuropathology section).

AD participants met criteria for dementia(27) or MCI(28) due to AD, or presented with a non-memory predominant AD phenotype as described below. All AD participants had a positive Aβ-PET scan.

Control (CTL) participants from the UCSF BrANCH study were clinically normal, functionally independent (CDR Global = 0), community-dwelling older adults. All CTL participants were Aβ-PET negative, lacked cognitive symptoms, and were free of major neurological, psychiatric, or medical conditions (e.g., sleep apnea, stroke, HIV) known to impact brain health. Absence of RHI in the CTL group was determined by completion of detailed self-report surveys of prior traumatic brain injury(29) and prior sport and military participation(30).

### Plasma proteomics

Venous blood was collected and stored in EDTA tubes (Alzheimer’s Disease Neuroimaging Initiative protocol) at − 80 °C until being packed with dry ice and sent to Olink (Olink Proteomics, Uppsala, Sweden) following standard shipping protocols. Plasma proteins were measured on the Olink Explore 3072 proximity extension assay (PEA)(31), which quantified 2,944 proteins. All samples were randomized by Olink prior to analysis on single plates. Relative quantification of protein abundance is reported as Normalized Protein eXpression (NPX) values in log2 scale. NPX values flagged by Olink with quality control warnings were removed from downstream analysis. Additional quality control of Olink data was performed using available code from a recent study examining the cross-platform validity of Olink against other proteomics platforms (SomaScan and TMT-mass spec)(26). Briefly, sample measurements that were below the limit of detection, defined as median log_2_ buffer signal plus 3 standard deviations (SD) of the assay’s buffer measurements (SD based on historically recorded background variance of the assay), were coded as missing. After removal of repeated measurements of the same UniProt protein assayed in different subpanels (N=18) and proteins with greater than 75% missingness (N=147 proteins), a total of 2779 proteins were retained for primary analysis. In addition to Olink, plasma samples were assayed for IL-6 (Meso Scale Discovery [MSD]), glial fibrillary acidic protein (GFAP; Quanterix Simoa), and neurofilament light chain (NfL; Quanterix Simoa), as previously described(19).

### Outlier removal and covariate adjustment

Outlier samples were identified (n=3 [2 CTL, 1 RHI]) and removed using a three-fold SD cutoff of Z-transformed sample connectivity, as previously described(32). Consistent with prior work(24, 32), nonparametric bootstrap regression was conducted to obtain a median estimated coefficient from 1,000 iterations of fitting for the effects of age and sex on each protein, while also modeling case status to protect against unwanted removal of case-related variance. Median estimated coefficients were subsequently multiplied by age and sex and subtracted from participant-specific protein values to derive adjusted protein values across the entire log_2_(abundance) matrix of cases and controls.

### Differential abundance analysis

Differential abundance analyses were performed on the post-processed log_2_(abundance) matrix. Differentially abundant proteins were identified using one-way ANOVA followed by Tukey post hoc correction of pairwise comparisons between RHI, AD, and CTL. Differential abundance was visualized using volcano plots with proteins color-coded by co-expression network module membership.

### Plasma protein co-expression network analysis

WGCNA(32, 33) was used to generate a plasma protein network from the *n*=2,779 log_2_ protein abundance × *n*=105 case–sample matrix that had undergone covariate correction and network connectivity outlier removal as described above. The WGCNA blockwiseModules function was run with the following parameters: power=7.5, deepSplit=4, minModuleSize=10, mergeCutHeight=0.07, TOMdenom=“mean”, bicor correlation, signed network type, PAM staging and PAM respects dendro as TRUE, with clustering completed within a single block. Module memberships were then iteratively reassigned to enforce kME table consistency, as previously described(32). Plasma protein co-expression network assignments were visualized as modules using publicly-available code that utilizes the igraph *R* package (https://github.com/edammer/netOps/tree/main).

### Gene ontology (GO) and cell type marker enrichment

GO enrichment of differential abundance lists and network modules was calculated as a Fisher exact test p value transformed to z score and visualized using the publicly-available GOparallel code (https://www.github.com/edammer/GOparallel), which pulls curated gene sets from the Bader Lab’s monthly update(34). As previously published(32), enrichment of network modules for cell type-specific marker gene symbols was performed using one-tailed Fisher’s test (https://github.com/edammer/CellTypeFET).

### Cognitive Assessment

A subset of cases underwent detailed neuropsychological assessments, as previously described(19). Global cognitive composite z-scores, representing a continuous index of cognitive severity, were calculated as the average of composite z-scores in the domains of episodic memory and executive functioning. Cognitive composites were residualized for the expected effects of age, sex, and education based on available data from the clinically normal BrANCH cohort (N ≥ 488 per composite). Correlations of protein co-expression modules and individual proteins with global cognition were performed specifically within RHI to identify proteomic signatures linked to cognitive function. One-tailed Fisher’s exact tests determined module-wise overrepresentation of proteins positively or negatively associated with global cognition, adjusting for FDR.

### Neuropathological Assessment

Participants that underwent autopsy (n=16) used standardized sampling and staining protocols in the UCSF Neurodegenerative Disease Brain Brank as described elsewhere(35, 36). Sampling procedures followed recommended guidelines for CTE, AD, frontotemporal lobar degeneration, and synucleinopathies classification(4, 37, 38) (phospho-tau antibody: CP-13, S202/T205, mouse, 1:250, courtesy of P. Davies; β-amyloid: 1-16, mouse, clone DE2, 1:500, Millipore; TDP-43 antibody: rabbit, 1:2000, Proteintech Group; alpha synuclein: LB509 mouse, 1:5000, courtesy of J. Trojanowski and V. Lee). AD burden (Aβ plaques and AD tau tangles) was defined as “None,” “Low,” “Moderate,” or “High” AD neuropathologic changes (ADNC) based on current NIA-AA criteria(38). CTE severity was defined according to McKee staging criteria(39) and as “High” or “Low” based on recently proposed classification methods that account for the number of brain regions with CTE-tau deposition (regardless of burden/density)(4). We first verified that participants with neuropathological diagnoses of CTE based on prior criteria also met recent consensus criteria (e.g., required neuronal tau inclusions)(3). We also reevaluated participants that were considered free of CTE pathology during initial autopsy evaluation by sampling additional regions (orbitofrontal cortex, superior/middle temporal cortex) for CTE tau pathology, per recent consensus group recommendations(4).

Burden of each proteinopathy (tau, TDP-43, beta-amyloid, alpha-synuclein) was calculated based on the sum of all semi-quantitative ratings (0 = none, 1 = mild/sparse, 2 = moderate, 3 = severe/frequent) in each sampled region for all inclusion types. Semi-quantitative ratings of neurodegeneration in each region were also summed based on observed neuron loss. We further evaluated individual proteinopathy burden and overall neurodegeneration within a medial temporal lobe region of interest (dentate gyrus, CA1/subiculum, CA2, CA3/CA4, entorhinal cortex, amygdala) based on prior research suggesting susceptibility of these structures to the neurodegenerative effects of RHI(3, 18).

### Other statistics

Statistical analyses were performed in R (v4.3.1). Boxplots represent the median, 25th, and 75th percentile extremes; thus, hinges of a box represent the interquartile range of the two middle quartiles of data within a group. Minimum and maximum data points define the extent of whiskers (error bars). Correlations were performed using Spearman’s rho (ρ) coefficients or biweight midcorrelations (bicor), as specified. Comparisons between two groups were performed by a two-sided t test. Comparisons among three or more groups were performed with ANOVA with Tukey’s pairwise comparison of significance. *P* values were adjusted for multiple comparisons by false discovery rate (FDR) correction according to the Benjamini-Hochberg method where indicated.

## Results

### Participant Characteristics

The primary study cohort included 22 RHI, 39 AD, and 44 CTL (**Table 1**). Participants with RHI were younger than the AD and CTL groups (RHI: 57.8±21.1, AD: 67.6±9.8, CTL: 70.6±7.2) and were all male (RHI: 100% males, AD: 44%, CTL: 64%). Both the RHI and CTL groups were less likely than the AD group to carry at least one copy of an *APOE e4* allele (RHI: 21%, CTL: 27%, AD: 60%). The RHI and AD groups did not differ in overall functional severity defined by CDR Sum of Boxes, although RHI did exhibit a lower proportion of patients with mild dementia compared to AD, defined by CDR-Global of 1 (RHI: 18%, AD: 46%).

**Table 1.**
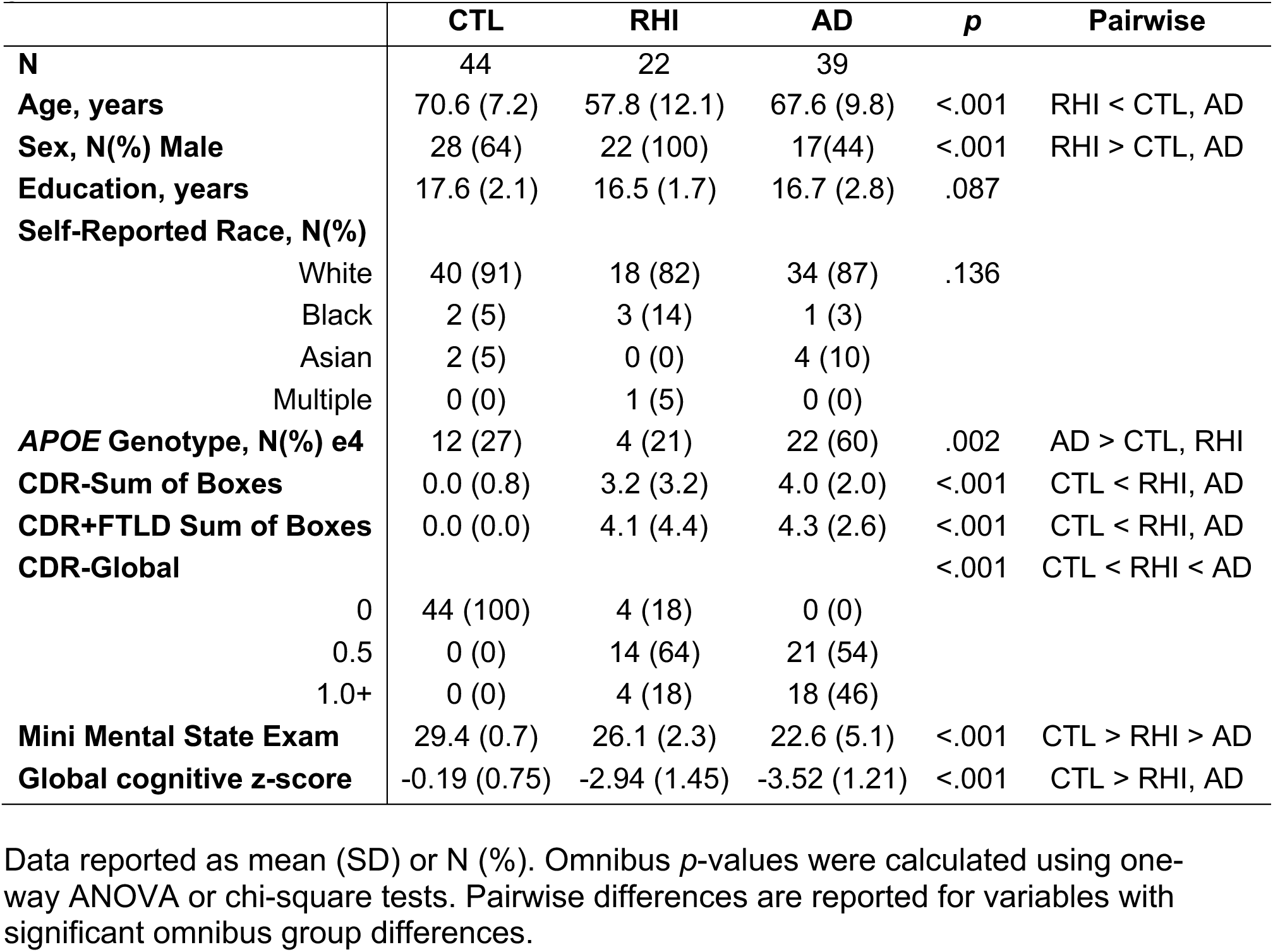
Descriptive demographic and clinical characteristics for the three primary study groups.

|  | **CTL** | **RHI** | **AD** | ***p*** | **Pairwise** |
| --- | --- | --- | --- | --- | --- |
| **N** | 44 | 22 | 39 |  |  |
| **Age, years** | 70.6 (7.2) | 57.8 (12.1) | 67.6 (9.8) | <.001 | RHI < CTL, AD |
| **Sex, N(%) Male** | 28 (64) | 22 (100) | 17(44) | <.001 | RHI > CTL, AD |
| **Education, years** | 17.6 (2.1) | 16.5 (1.7) | 16.7 (2.8) | .087 |  |
| **Self-Reported Race, N(%)** |  |  |  |  |  |
| White | 40 (91) | 18 (82) | 34 (87) | .136 |  |
| Black | 2 (5) | 3 (14) | 1 (3) |  |  |
| Asian | 2 (5) | 0 (0) | 4 (10) |  |  |
| Multiple | 0 (0) | 1 (5) | 0 (0) |  |  |
| **APOE Genotype, N(%) e4** | 12 (27) | 4 (21) | 22 (60) | .002 | AD > CTL, RHI |
| **CDR-Sum of Boxes** | 0.0 (0.8) | 3.2 (3.2) | 4.0 (2.0) | <.001 | CTL < RHI, AD |
| **CDR+FTLD Sum of Boxes** | 0.0 (0.0) | 4.1 (4.4) | 4.3 (2.6) | <.001 | CTL < RHI, AD |
| **CDR-Global** |  |  |  | <.001 | CTL < RHI < AD |
| 0 | 44 (100) | 4 (18) | 0 (0) |  |  |
| 0.5 | 0 (0) | 14 (64) | 21 (54) |  |  |
| 1.0+ | 0 (0) | 4 (18) | 18 (46) |  |  |
| **Mini Mental State Exam** | 29.4 (0.7) | 26.1 (2.3) | 22.6 (5.1) | <.001 | CTL > RHI > AD |
| **Global cognitive z-score** | -0.19 (0.75) | -2.94 (1.45) | -3.52 (1.21) | <.001 | CTL > RHI, AD |

Data reported as mean (SD) or N (%). Omnibus *p*-values were calculated using one-way ANOVA or chi-square tests. Pairwise differences are reported for variables with significant omnibus group differences.

Of the 22 participants with RHI, 18 (82%) met research criteria for TES (N = 6 Probable TES, N = 7 Possible TES, N = 5 Suggestive of TES). Two participants with RHI were felt to not meet TES criteria due to a clinical phenotype better explained by an alternative diagnosis (behavioral variant frontotemporal dementia), one did not show clear evidence of progressive decline over time, and one had insufficient RHI exposure duration data available to determine TES status.

### Differential Abundance

A total of 2,779 proteins were included in primary analyses after filtering out proteins flagged for QC warnings and proteins with high missingness. We performed differential abundance analysis to identify proteins that were significantly altered in plasma between RHI compared to AD or CTL samples (Supplementary Table 1). We observed a striking pattern of increased protein abundance in RHI vs. CTL (Fig. 1A-B). After Tukey’s post-hoc test (Tukey *p* <0.05), 434 proteins were significantly increased in RHI vs. CTL whereas only 6 proteins were decreased in RHI vs. CTL. Pathway analysis of proteins increased in RHI vs. CTL (Fig. 1C) revealed strong enrichment for mitochondrial proteins, including top hits MECR, GATD3, KIFBP, BECN1, and TOMM20, as well as enrichment for microtubule cytoskeleton (RILP, CD2AP) and protein serine/threonine phosphatase complex proteins (PPP3R1, PPP2R5A, PPP1CC). RHI vs. AD comparisons revealed a similar shift towards increased protein abundance in RHI, with 193 proteins increased and 14 decreased in RHI (Fig. 1B). Pathway analysis of proteins increased in RHI vs. AD (Fig. 1C) revealed immune pathway enrichment, including TNF-receptor superfamily proteins (TNFRSF1B, TNFRSF1A, TNFRSF11A, TNFRSF14), as well as mitochondrial and phosphatase complex proteins that overlapped with RHI vs. CTL results (MECR, TOMM20, PPP3R1).

**Figure 1.**
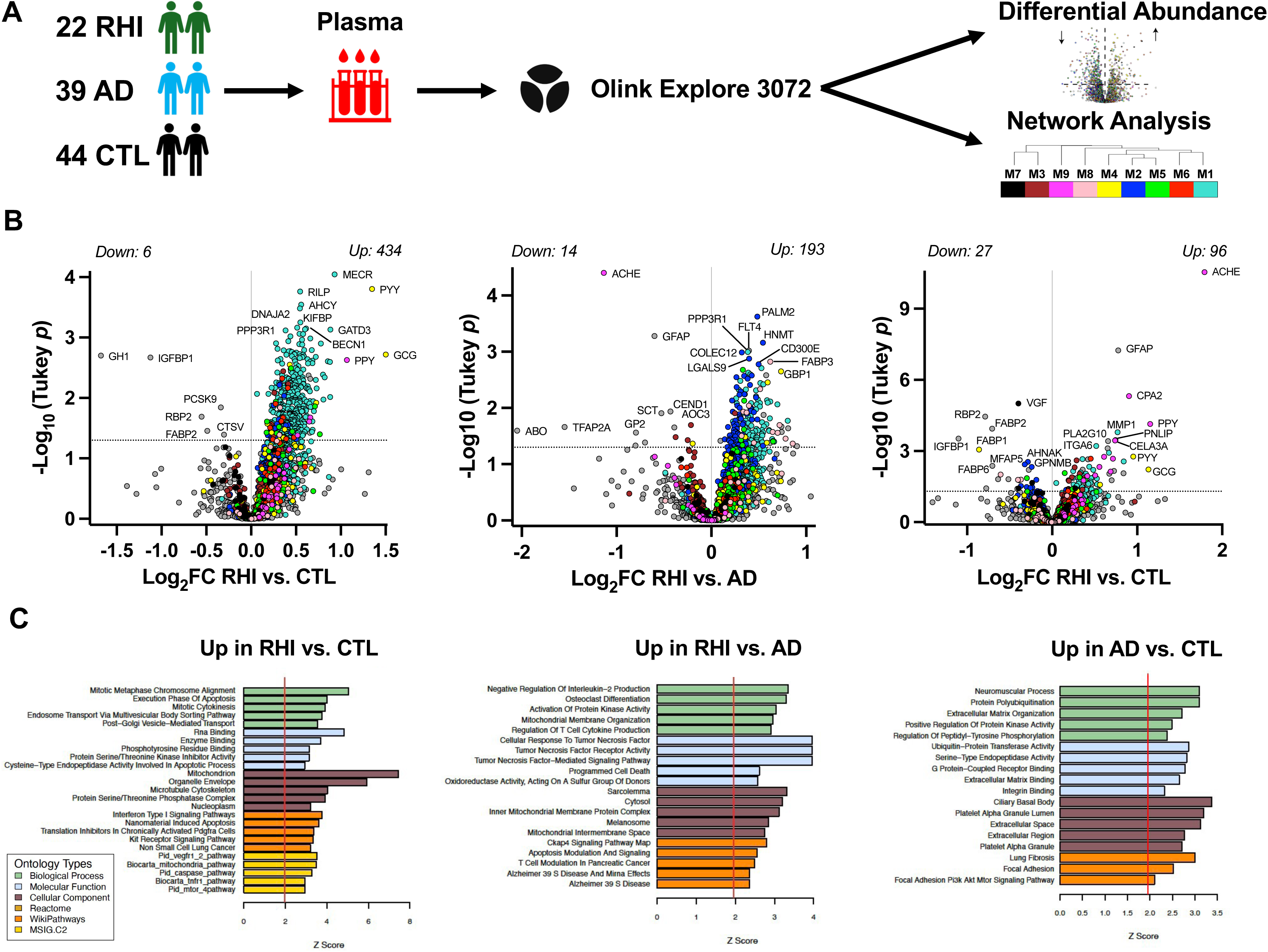
Study design and differential plasma protein abundance across RHI, AD, and control (CTL) Legend: **A)** Plasma was collected in a clinical cohort of 22 AD-biomarker negative patients with substantial prior repetitive head injury (RHI), 39 AD-biomarker positive patients with cognitive impairment, and 44 AD-biomarker negative controls (CTL). Plasma proteomics was performed on the Olink Explore proximity-extension assay. After data processing, a total of 2,779 proteins were examined in downstream differential abundance and weighted gene co-expression network analysis (WGCNA). **B)** Differential abundance analyses examined individual plasma protein differences across RHI, AD, and CTL. Pairwise differential protein abundance is represented by volcano plots of average log_2_ difference by negative log_10_ *p*-value for each given comparison. Proteins are colored by the network module in which they were assigned to. Pairwise comparisons were performed using one-way ANOVA with Tukey post-hoc test. The horizontal dotted line represents Tukey *p* < 0.05. **C)** Gene ontology (GO) analysis was performed on proteins exhibiting increased abundance in RHI vs. CTL, RHI vs. AD, and AD vs. CTL. Pathway enrichment is shown by *z* score, transformed from a Fisher’s exact test.

Supporting the validity of findings from this multiplex, ACHE and GFAP, two proteins consistently linked to AD(40–42), were the top two most significantly increased proteins in AD compared to both RHI and CTL (Fig. 1B). Additional AD vs. CTL hits (96 increased, 27 decreased) included decreased abundance of fatty acid binding proteins (FABP1, FABP2, FABP6) and neuropeptide VGF, as well as increased abundance of lipid enzymes (PNLIP, PLA2G10) and extracellular matrix-associated proteins (ITGA6, MMP1).

### Network Analysis

We leveraged WGCNA to identify communities of proteins that are co-expressed across plasma samples and dysregulated in RHI. WGCNA revealed nine protein co-expression modules (Fig. 2A). Protein module membership assignments are provided in Supplementary Table 2 and network module protein graphs are visualized in Supplementary Fig. 1. Pathway and cell type enrichment analyses were performed to identify the primary ontology used for module annotation (Supplementary Table 3; Supplementary Fig. 2). Seven out of the nine network module eigenproteins were elevated in RHI compared to CTL and/or AD (Tukey *p* < 0.05), recapitulating the overall shift towards increased protein abundance that was observed in volcano plots (Fig. 2B, Supplementary Table 4). M1 intracellular signaling, enriched for ontology terms related to processes of cell division, cytoskeletal organization, and mitochondrial function, was the largest module (n=853) and harbored the highest proportion of differentially abundant proteins in RHI vs. CTL (43% of the module; Fig. 2C). M2 TNF-signaling/cell adhesion, enriched for microglial cell type markers and extracellular matrix/immune-related ontology terms, was elevated in RHI vs. AD and harbored TNF-receptor superfamily proteins identified in differential abundance analysis. M2 clustered alongside M4 cell metabolism and M5 chemotaxis, two additional modules that were elevated in RHI and also enriched for extracellular and immunometabolic processes. M9 digestion and M6 cellular detoxification were both significantly elevated in RHI and AD compared to controls.

**Figure 2.**
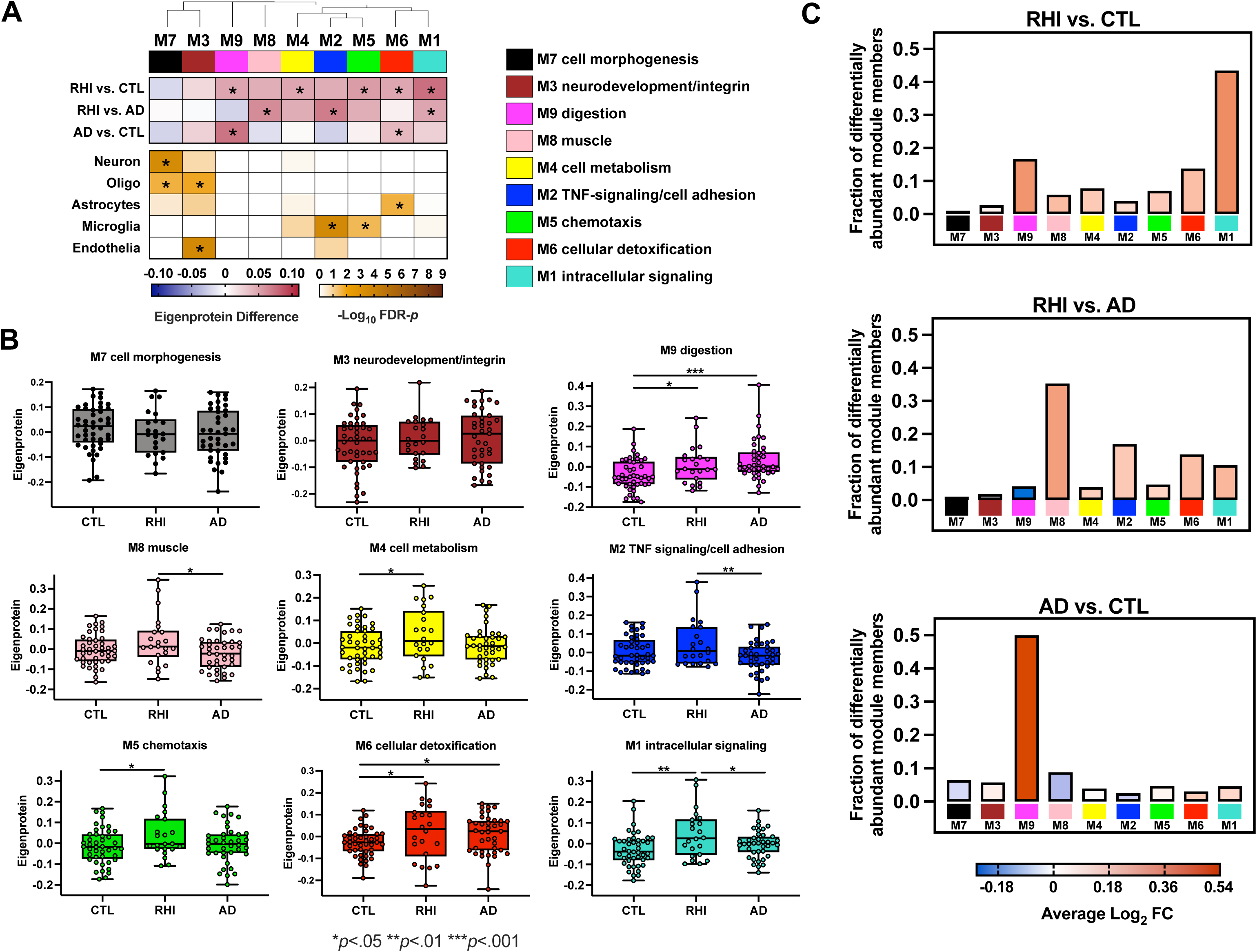
Plasma protein co-expression network across RHI, AD, and CTL. Legend: **A)** A plasma protein co-expression network was built using weighted gene correlational network analysis (WGCNA). The plasma network consisted of 9 protein co-expression modules. Module relatedness is shown in the dendrogram above the heatmap. GO analysis was used to identify the principal biology represented by each module. Increased eigenprotein abundance for each comparison is indicated in red, whereas decreased eigenprotein abundance is indicated in blue. The cell type nature of each module was assessed by module protein overlap with cell-type-specific marker lists of neurons, oligodendrocytes, astrocytes, microglia and endothelia. Asterisks denote statistically significant comparisons or enrichment. **B)** Module eigenprotein levels by case status across the 9 plasma network modules. Box plots represent the median and 25th and 75th percentiles, and box hinges represent the interquartile range of the two middle quartiles within a group. Min and max data points define the extent of whiskers (error bars). \*\*\**p*<.001, \*\**p*<.01, \**p*<.05. **C)** Bar heights represent the fraction of module member proteins that exhibited differential abundance. Bars are color coded by the average log_2_ fold-change (FC) of module member proteins.

### Orthogonal plasma biomarker measurement highlights IL-6 as a marker of RHI-related proteomic dysregulation

We have previously shown that plasma concentration of IL-6 is elevated in amyloid-negative RHI compared to controls and AD(19). We examined correlations between Olink-based proteomic signatures and orthogonally measured plasma immunoassays for IL-6 (MSD) and GFAP and NfL (Quanterix Simoa) to determine the degree to which RHI plasma proteomic signatures converged with each more established plasma biomarker. Cross-platform correlations were large when examining Olink and orthogonally-measured IL-6 (*r*=0.91), GFAP (*r*=0.75), and NfL/NEFL (*r*=0.86) across the study cohort (Fig. 3A). M5 chemotaxis, which included Olink-based IL-6 as a module member, demonstrated the strongest correlations with MSD-based IL-6 in RHI, AD, and CTL (Spearman’s ρ range: 0.46 to 0.65; Supplementary Table 5; Fig. 3B). The remaining significant correlations between modules and IL-6 were only observed among modules that were elevated in RHI (M1, M2, M4; Fig. 3C). Notably, none of the plasma modules exhibited significant correlations with GFAP or NfL in RHI cases, whereas M7 cell morphogenesis, M3 neurodevelopment/integrin, and M2 TNF-signaling/cell adhesion significantly correlated with GFAP (M7, M3, M2) and/or NfL (M7, M2) in AD (ρs > 0.35). These results validate individual and network-based Olink plasma biomarkers against targeted immunoassays and further support IL-6 as a sensitive marker of RHI-related pathophysiology, particularly in comparison to canonical plasma biomarkers of neurodegeneration.

**Figure 3.**
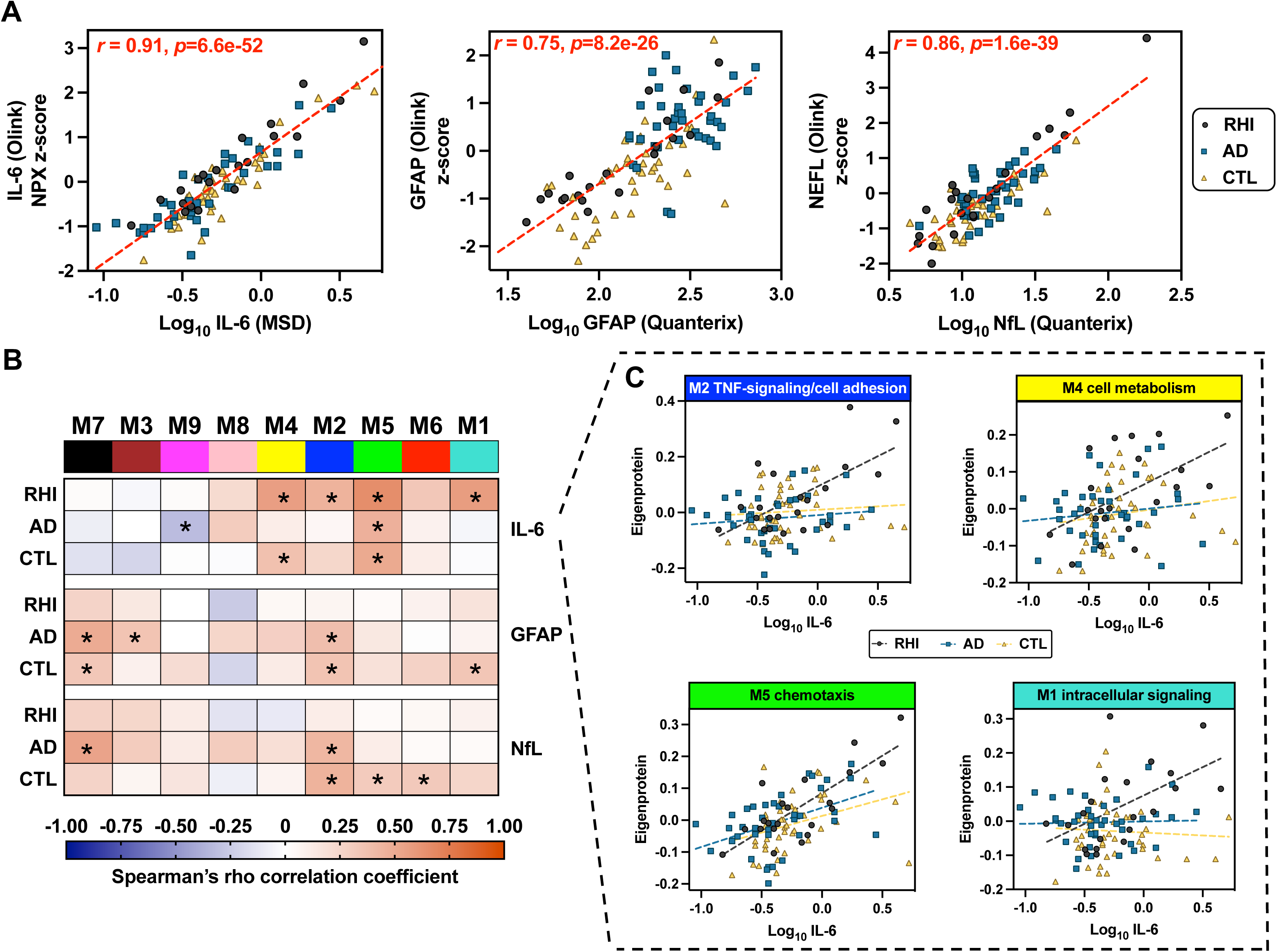
Olink-based plasma module associations with orthogonally-measured IL-6, GFAP, and NFL. Legend: **A)** Correlation of IL-6, GFAP, and NFL measured by immunoassay (x-axis) to Olink-based IL-6, GFAP, and NFL (y-axis). MSD = Meso Scale Discovery. **B)** Heatmap of plasma module eigenprotein correlations (Spearman’s rho) with immunoassay-based IL-6, GFAP, and NFL, stratified by study group. Asterisks denote statistically significant correlations. **C)** Scatterplot of select module eigenprotein correlations with IL-6 across RHI, AD, and CTL.

### RHI plasma protein co-expression signatures are reflected among cases with pathology-confirmed CTE and correlate with neuropathology burden

RHI is largely a prerequisite for developing CTE pathology(43), yet RHI alone does not guarantee the presence of CTE. To determine whether the protein abundance patterns observed in our RHI cohort may be related to underlying CTE, we compared individual protein and co-expression module abundances between RHI individuals with autopsy-confirmed CTE and non-RHI cases that were CTE-negative at autopsy (Supplementary Table 6). CTE+ cases (n=7) included 5 amyloid-negative RHI that were included in the primary RHI grouping as well as two additional CTE+ cases that were excluded from primary analyses due to amyloid-positivity (Fig. 3). CTE-negative cases (n=9) included 5 AD patients and 4 CTL from primary analyses. Similar to RHI results, CTE+ cases exhibited a shift towards increased protein abundance vs. CTE-negative (290 proteins increased, 10 proteins decreased at nominal *p* < 0.05; Supplementary Table 7; Fig. 4A, B). Proteins elevated in CTE were enriched for ontology terms related to immune (“acute-phase response”, “cellular response to IL-7”), metabolic/antioxidant (“glutathione metabolic process”), cytoskeleton, and cell division processes (Fig. 4C). All 7 co-expression modules that were increased in RHI exhibited elevations of a similar or higher magnitude in CTE+ vs. CTE-cases (Supplementary Table 8). Of those 7, only four reached statistical significance (*p* < 0.05) likely related to the smaller analytic sample size: M1 intracellular signaling, M2 TNF-signaling/cell adhesion, M4 cell metabolism, and M9 digestion (Fig. 4D). Of the 80 total proteins that were differentially abundant in both RHI (vs. CTL) and CTE+ (vs. CTE-), 69 (88%) were assigned to M1, including mitochondrial (GATD3, ACOT13, NFU1) and cell division proteins (PPP2R5A, UFD1) that exhibited high log-fold changes (>0.69) in RHI.

**Figure 4.**
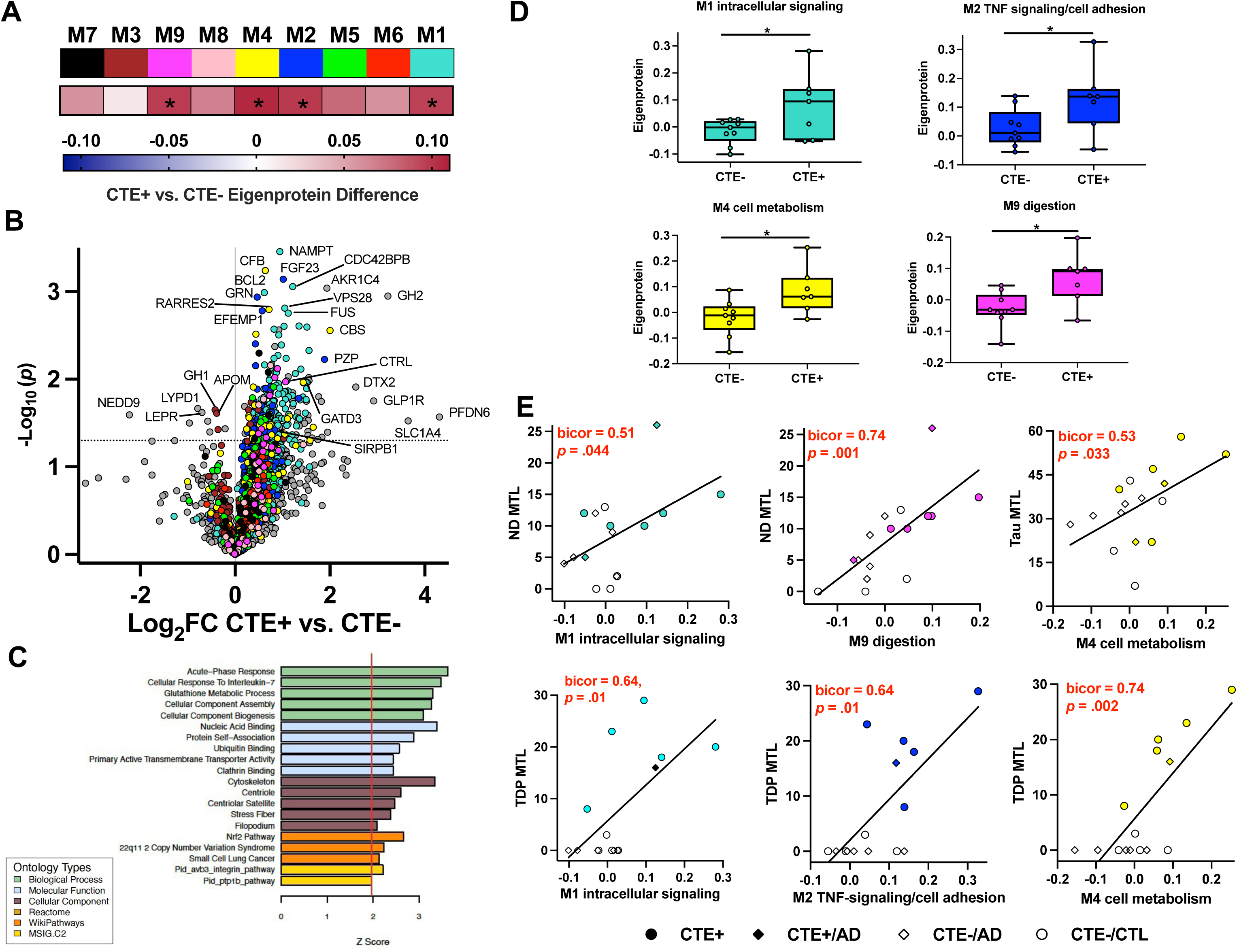
Plasma proteome associations with pathology-confirmed chronic traumatic encephalopathy (CTE) and neuropathological burden Legend: **A)** Heatmap of plasma module eigenprotein differences between CTE+ and CTE-cases. Asterisks denote statistically significant differences. **B)** Differential abundance of plasma proteins between CTE+ and CTE-. Differential protein abundance is represented by a volcano plot of average log_2_ difference by negative log_10_ *p*-value. Proteins are colored by the module in which they were assigned to. The horizontal dotted line represents *p* < 0.05. **C)** GO analysis was performed on proteins exhibiting increased abundance in CTE+ vs. CTE-. Pathway enrichment is shown by *z* score, transformed from a Fisher’s exact test. **D)** Eigenprotein levels by CTE+ and CTE-for the 4 plasma network modules with significant differences. Box plots represent the median and 25th and 75th percentiles, and box hinges represent the interquartile range of the two middle quartiles within a group. Min and max data points define the extent of whiskers (error bars). **E)** Scatterplot of select module eigenprotein correlations (biweight midcorrelations [bicor]) with semi-quantitative ratings of neurodegeneration (ND), tau, and TDP-43. Data points are color and symbol coded based on CTE status with and without comorbid AD neuropathology.

We next evaluated how well the four co-expression modules significantly elevated in CTE correlated with semi-quantitative ratings of neurodegeneration and neuropathology burden in all sampled regions and in an MTL region of interest (Supplementary Table 8; Fig. 4E). Among the overall autopsy cohort, higher eigenproteins in three of the four modules associated with greater TDP-43 proteinopathy burden in the MTL: M1 intracellular signaling (bicor = 0.64, p = .01), M2 TNF-signaling/cell adhesion (bicor = 0.64, p = .01), M4 cell metabolism (bicor = 0.74, p=.002). M1 intracellular signaling (bicor = 0.51, p = .044) and M9 digestion (bicor = 0.74, p = .001) were associated with greater MTL neurodegeneration, and M4 cell metabolism was associated with greater tau proteinopathy burden in the MTL (bicor = 0.53, p = .033). No clear associations emerged with alpha-synuclein or beta-amyloid burden (Supplementary Table 8).

Overall, these results suggest that plasma protein elevations observed in RHI track with the diverse neuropathological changes observed in autopsy-confirmed CTE cases.

### M2 TNF-signaling/cell adhesion tracks with cognitive function in RHI

Differential abundance of plasma proteins in symptomatic RHI/CTE cases could reflect pathophysiological processes that start early in the disease course and increase with clinical symptom progression, or emerge only at later stages of clinical impairment. We performed differential correlations with global cognitive scores to identify plasma protein signatures sensitive to RHI cognitive severity. Out of the nine co-expression modules, M2 TNF-signaling/cell adhesion (ρ=-0.56, *p*=0.024) and M3 neurodevelopment/integrin (ρ=-0.64, *p*=0.008) exhibited significant, negative correlations with global cognition in RHI (Supplementary Table 9; Fig. 5C, D). Differential correlation analysis with individual proteins confirmed overlap with the associations observed between M2 and M3 with global cognition in RHI (Supplementary Table 10; Fig. 5A, B); that is, both modules had significant overrepresentation of individual proteins negatively associated with global cognition (M2 FDR-*p*=4.8e-17, M3 FDR-*p=*0.001). Of the 147 proteins nominally-associated with global cognition, 24 also differed between RHI vs. CTL and/or CTE+ vs. CTE-, including 11 proteins from M2 (Fig. 5E) linked to adaptive immunity (e.g., CD72, FCAMR, TNFSF13B) and cell adhesion/growth factor signaling (e.g., TGFBR2, FBLN2, EFEMP1). GATD3 and PPP3R1 were top M1 intracellular signaling hits from differential abundance analyses that were also negatively correlated with global cognition. These findings collectively highlight extracellular proteins, particularly M2 proteins involved in immunovascular signaling, as markers of RHI-related cognitive changes that may track with symptom progression throughout the disease course.

**Figure 5.**
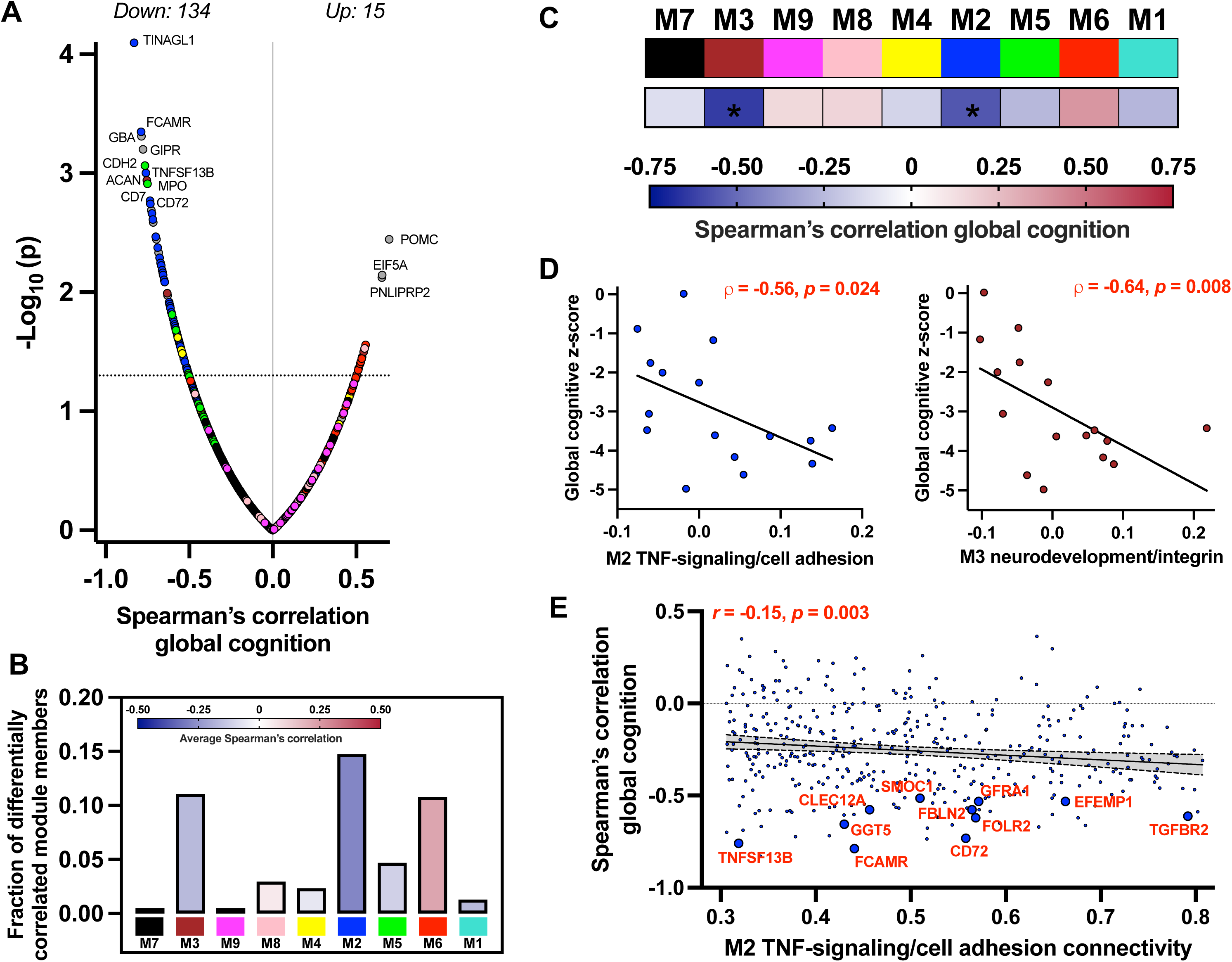
Plasma proteome associations with global cognition in RHI cases. Legend: **A)** Differential correlations (Spearman’s rho) of plasma proteins with global cognitive z-scores in RHI cases. Differential correlation is represented by a volcano plot of Spearman’s rho correlations by negative log_10_ *p*-value. Proteins are colored by the module in which they were assigned to. The horizontal dotted line represents *p* < 0.05. **B)** Bar heights represent the fraction of module member proteins that exhibited significant correlation with global cognition. Bars are color coded by the average correlation coefficient of module member proteins. **C)** Heatmap of plasma module eigenprotein correlations with global cognition in RHI cases. Asterisks denote statistically significant correlations. **D)** Scatterplot of statistically significant module eigenprotein correlations with global cognitive z-scores (M2 TNF-signaling/cell adhesion and M3 neurodevelopment/integrin). **E)** For proteins assigned to M2 TNF-signaling/cell adhesion, an individual protein’s strength of connectivity to M2 (x-axis) is plotted against the individual protein’s correlation with global cognitive z-score (y-axis). Proteins that exhibited stronger intramodular connectivity tended to exhibit stronger relationships with global cognition. Annotated proteins are M2 members that were significantly negatively correlated with global cognition and exhibited increased abundance in RHI vs. CTL and/or CTE+ vs. CTE-comparisons.

## DISCUSSION

We used quantitative plasma proteomics in cognitively impaired older adults with prior RHI to interrogate the disrupted molecular pathways that may underlie later-life onset of RHI-related neurodegenerative changes. Compared to healthy controls and individuals with AD, we observed a large shift toward increased plasma protein abundance in RHI. The most upregulated pathways involved proteins associated with mitochondrial function, metabolism, and immune activation. Network analysis leveraging multivariate protein co-expression revealed a diverse set of protein communities elevated in RHI, including M1 intracellular signaling and M2 TNF-signaling/cell adhesion. Network derived pathways dysregulated in RHI were 1) similarly dysregulated in the subset of participants with autopsy-confirmed CTE, 2) correlated with greater neuropathological burden in the medial temporal lobe (tau and TDP43 proteinopathy), 3) selectively correlated with orthogonally-assayed IL-6, and 4) linked to worse global cognition in RHI. Overall, our findings provide a comprehensive index of plasma proteins that correlate with clinical severity in RHI and pathological severity in CTE. These data will support targeted and testable hypotheses about blood-detectable molecular pathways linking RHI exposure to neurodegenerative disease.

M1 intracellular signaling was the largest module (>800 proteins) and harbored the greatest proportion of differentially abundant proteins in RHI versus controls. The strong M1 eigenprotein elevations in RHI compared to controls and AD suggest the presence of large-scale changes in intracellular signaling pathways linked to mitochondrial function, calcium homeostasis, cell division, and cytoskeletal integrity in RHI. Human iPSC models demonstrate marked bioenergetic failures and mitochondrial dysfunction in tauopathies(44, 45) and recent proteomic studies report mitochondrial and cytoskeletal protein abundance changes in brain and CSF across multiple tauopathies(25, 46). This supports the observed effects in an RHI cohort putatively at higher risk for CTE but also often possessing other neuropathologic changes like TDP-43 proteinopathy. Notably, M1 contained proteins such as MECR, TOMM40, and GATD3, which are critical for mitochondrial protein synthesis and transport, and PPP3R1, a protein involved in RNA transcription and excitatory postsynaptic signaling that also harbors a genetic risk loci for AD(47).

We have previously demonstrated peripheral immune dysregulation in a cohort of older adults with RHI using targeted measurement of plasma IL-6 (19). Here we extend observations of RHI-related immune dysregulation with large-scale plasma proteomics. The M2 module, enriched in TNF-super family proteins, extracellular matrix constituents, immunoglobulin-binding proteins, and microglial cell type markers, was elevated in RHI and CTE cases and harbored proteins that correlated with global cognitive function. Mass-spectrometry proteomics performed in CTE brain tissue has previously identified multiple elevated glial cell type-linked modules, implicating brain inflammation as a key component of CTE pathogenesis(24). Interestingly, a prior description of the CTE brain tissue proteome noted minimal differences between early CTE (Stage I-II) and controls, with two exceptions – one module enriched for extracellular matrix proteins, and a second module enriched with immunoglobulins that uniquely and sharply increased with CTE Stage II(24). Collectively, there is converging evidence across brain and blood proteomics for an upstream role of extracellular matrix degradation and immune activation early in RHI-related disease pathogenesis.

RHI is not a neurodegenerative disease but rather an environmental exposure variable that increases risk for CTE and other neurodegenerative diseases. The high frequency and different combinations of CTE co-pathology reported in RHI case series(2, 3, 7) suggest RHI is a common upstream exposure that can increase risk for diseases beyond CTE. Plasma proteomic signatures in our living RHI cohort were recapitulated in a subset of seven brain donors with autopsy-confirmed CTE, where M1, M2, M4, and M9 were all significantly elevated compared to nine brain donors without CTE. Brain donors with CTE all had mixed neuropathology and included two individuals with severe AD. M1 intracellular signaling and M2 TNF-signaling/cell adhesion correlated with medial temporal lobe TDP43 burden, while M4 cell metabolism correlated with both TDP-43 and tau burden. Larger clinico-pathological studies including brain donors with relatively “pure” CTE as well as mechanistic preclinical studies will be necessary for determining whether observed alterations in the plasma proteome are related to CTE versus other pathologies. Regardless, alteration in these immune and metabolic modules may offer a unifying framework that ties RHI to neurodegeneration. Further, module hub proteins may inform targets for risk screening and biomarker or therapeutic development.

Lack of in vivo CTE biomarkers and the broad spectrum of neuropathological consequences of RHI beyond CTE complicate disease-specific attributions for altered molecular pathways or clinical symptoms. For example, medial temporal lobe atrophy and memory loss are common in cognitively impaired older adults with RHI, but corresponding neuropathological findings are highly variable(3, 11, 18–20). Neurobehavioral dysregulation (e.g., explosivity, emotional lability, aggression) is commonly reported in individuals with RHI but without an established neuropathological correlate(48). Anecdotally, individuals with extensive prior RHI such as former professional American football players often present in clinics with severe complaints and distress disproportionate to objective cognitive test scores or brain MRI, which often evidence minimal to mild dysfunction. Employing discovery-based plasma proteomics sheds new light on alternative pathophysiological contributors (e.g., immune or metabolic dysfunction), and thus therapeutic targets, for the debilitating symptoms reported by many individuals with prior RHI(49).

Collectively, there is important knowledge gained from identifying and characterizing protein co-expression modules implicated in the downstream effects of RHI, but these modules reflect broad and complex biological functions. The ultimate impact of these preliminary findings rests on continued analysis of individual proteins within the modules, replication in independent cohorts, and potentially a bedside-to-bench approach to test causal inferences related to RHI in a controlled environment. Different proteomics modalities beyond plasma are also essential. The degree to which our study of peripheral protein expression measured in plasma reflects central nervous system dysfunction is unclear. Nonetheless, the bidirectional influence of blood-and-brain pathophysiology is well established in the context of neurodegenerative diseases, particularly for inflammation and immune-linked molecular pathways(50–53).

There are several study limitations. We cannot infer causality between RHI and observed group differences in protein expressions, and most RHI participants did not undergo autopsy, so firm conclusions cannot be drawn regarding presence or combination of neurodegenerative changes. The RHI group was defined by a common environmental exposure history, but the subset who underwent autopsy suggested diverse neuropathology including some with conditions known to have severe inflammatory and neurodegenerative effects (e.g., ALS). Results are not necessarily CTE-specific, even among those with autopsy-confirmed CTE, but may still have relevance for RHI given high rates of TDP-43 co-pathology in RHI and CTE. Neuropathological analyses involved overall and medial temporal lobe-specific semi-quantitative burden ratings. Investigating other brain regions and neuropathological changes (e.g., gliosis, neuroinflammation) and incorporating more sophisticated quantitative assessment methods are important next steps in validating findings. While RHI participants underwent extensive workup including amyloid PET imaging to rule out contributions of AD pathology, the sample was relatively small, and we did not have an independent validation cohort. Replication of findings in other RHI populations including different types and extent of exposure is critical. The RHI participants were enrolled at a memory and aging research center with concerns for their cognitive and/or behavioral function, which is not representative of all individuals who have had prior RHI. The study sample lacked ethno-racial diversity and the RHI group was all males. Findings may not generalize to all populations given the likely complex interaction of biological sex, social determinants of health, genetic, and epigenetic factors relevant to plasma proteomic alterations and RHI. Findings came from plasma without cerebrospinal fluid or brain tissue proteomics, which limits our understanding of central nervous system specificity.

## CONCLUSIONS

Individuals with prior repetitive head trauma have plasma proteome alterations distinct from healthy controls and individuals with Alzheimer’s disease. Implicated molecular pathways in plasma were tied to immune response, mitochondrial function, and cell metabolism; correlated with TDP-43 and tau neuropathological burden at autopsy; and were associated with worse global cognition. Individual differentially expressed proteins and protein co-expression modules may inform mechanisms linking RHI to increased dementia risk, thus guiding diagnostic biomarker and therapeutic development for at-risk populations.

## List of Abbreviations

AD: Alzheimer’s disease
*APOE*: Apolipoprotein E
CDR: Clinical Dementia Rating scale
CTE: Chronic traumatic encephalopathy
CTL: Controls
GFAP: Glial fibrillary acidic protein
GO: Gene ontology
IL6: Interleukin-6
NfL/NEFL: Neurofilament light chain
PET: Positron emission tomography
RHI: Repetitive head impacts
TDP43: Transactive response DNA binding protein of 43 kDa
TES: Traumatic encephalopathy syndrome
WGCNA: Weighted gene co-expression network analysis

## DECLARATIONS

### Ethics approval and consent to participate

All data were collected following study procedures that were reviewed and approved by the UCSF institutional review board (IRB-01) and participants provided informed consent prior to participation.

### Consent for publication

Not Applicable

### Availability of Data and Materials

All study data are available on reasonable request made to the UCSF Memory and Aging Center. Academic, not-for-profit investigators can request data for professional education and for research studies. Requests can be made online (https://memory.ucsf.edu/research-trials/professional/open-science). Datasets used for the analyses for the current study are also available from the corresponding author on reasonable request.

### Competing interests

JHK has provided consultation to Biogen. GDR receives research funding from Avid Radiopharmaceuticals, GE Healthcare, Genentech, Life Molecular Imaging, and has served as consultant for Alector, Eli Lilly, Genentech, GE Healthcare, Roche, Johnson & Johnson, and Merck, and is an Associate Editor for JAMA Neurology. SS has provided consultation to Techspert.io and Putnam Associates.

### Funding

We thank the following funding sources who have supported our work: NIH ADRC (P30AG066506; MPI: Smith, Duara, Loewenstein) and 1Florida ADRC Development Grant to BMA; Alzheimer’s Association (AARF-23-1145318), New Vision Research (CCAD2024-001-1) and American Academy of Neurology Scholarship to RS; NIH ADRC (P30AG062422) to GDR; NIH (P01AG019724) to BLM; NIH (R01(s) AG032289 and AG048234) and Larry L. Hillblom Network Grant (2014-A-004-NET) to JHK; NIH (R01AG072475) to KBC.

### Author Contributions

Study conception and design: RS, BMA

Data acquisition, collection, and analysis: RS, SS, LTG, WWS, BLM, GDR, JHK, BMA

Manuscript drafting: RS, BMA

Manuscript review for scientific content: KBC, SR, PC, JFA, SS, LTG, WWS, BLM, GDR, JHK

Final approval of manuscript: All authors

## Acknowledgements

We are deeply grateful to the study participants and their families for participating in the UCSF Memory and Aging Center research programs. We additionally thank the teams of coordinators and administrative staffs who have provided integral support for the ongoing execution of research at our centers.

## Supplementary Table Titles

**Supplementary Table 1.** Plasma protein differential abundance across RHI, AD, and CTL. **Supplementary Table 2.** Plasma network module membership. **Supplementary Table 3.** Plasma network module gene ontology (GO) enrichment. **Supplementary Table 4.** Plasma network module eigenprotein differences across RHI, AD, and CTL. **Supplementary Table 5.** Plasma network module correlations with immunoassay-based IL-6, GFAP, and NFL. **Supplementary Table 6.** Autopsy subcohort characteristics. **Supplementary Table 7.** Plasma protein differential abundance between CTE+ and CTE-. **Supplementary Table 8.** Plasma network module associations with CTE and neuropathology ratings. **Supplementary Table 9.** Plasma network module associations with global cognition in RHI. **Supplementary Table 10.** Plasma protein differential correlations with global cognition in RHI.

## Supplementary Figure Legends

**Supplementary Figure 1.**

Pathway analysis on plasma protein co-expression network modules. Gene ontology (GO) analysis was performed to ascertain the principal biology represented by the constituent proteins in each module. Enrichment for a given ontology is shown by *z* score, transformed from a Fisher’s exact test.

**Supplementary Figure 2.**

Plasma network module protein graphs. The size of each circle indicates the relative eigenprotein correlation value (kME) in each network module. Those proteins with the largest kME are considered “hub” proteins within the module, and explain the largest variance in module expression. The top 150 proteins by kME for each module are shown.

