## Supplementary Figure 1 for "Interrogating the plasma proteome of repetitive head impact exposure and chronic traumatic encephalopathy"

**M7 black module: cell morphogenesis**

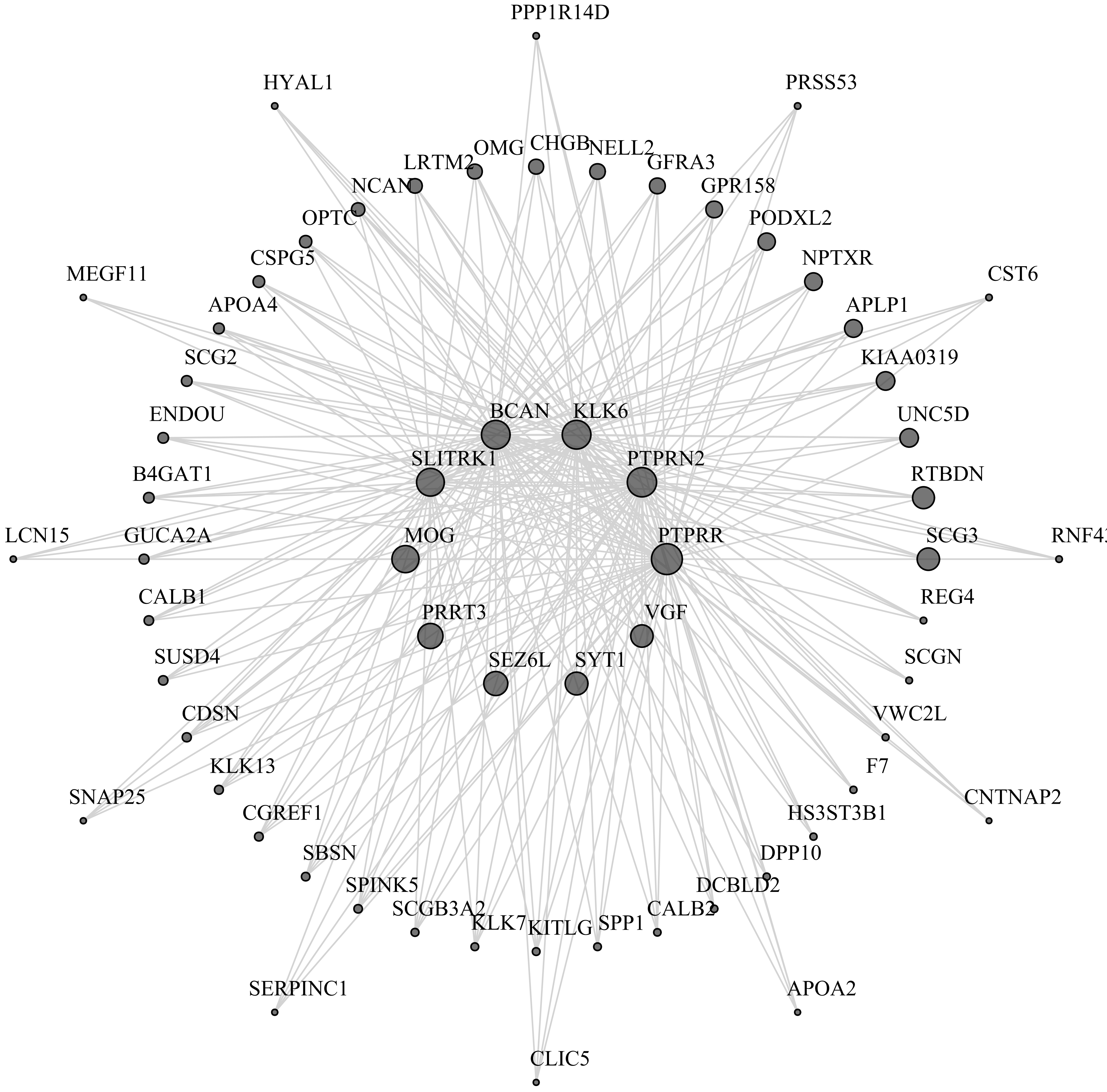

**M3 brown module: neurodevelopment/integrin**

**Supplementary Figure 1**

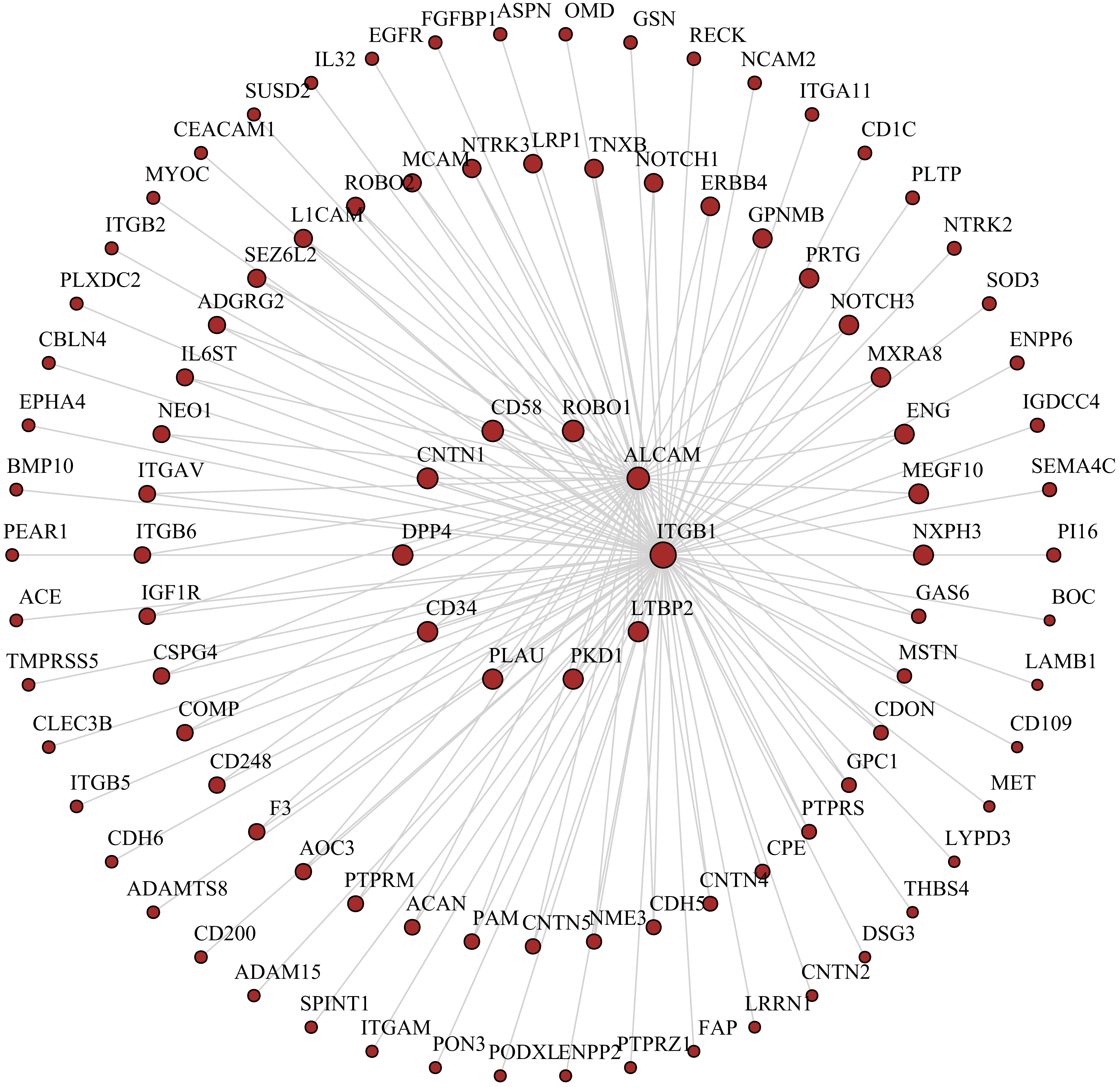

M9 magenta module: digestion

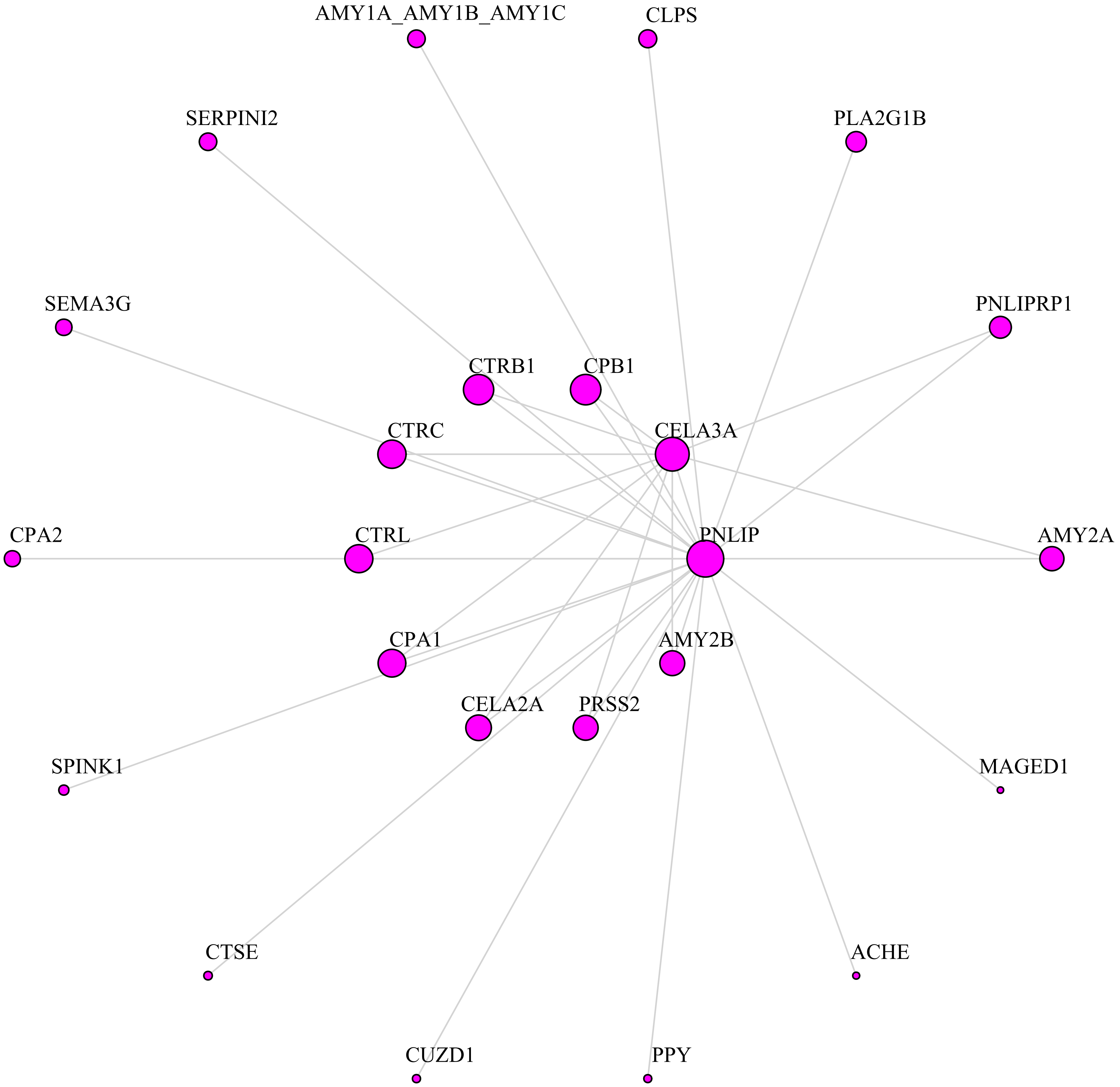

Supplementary Figure 1

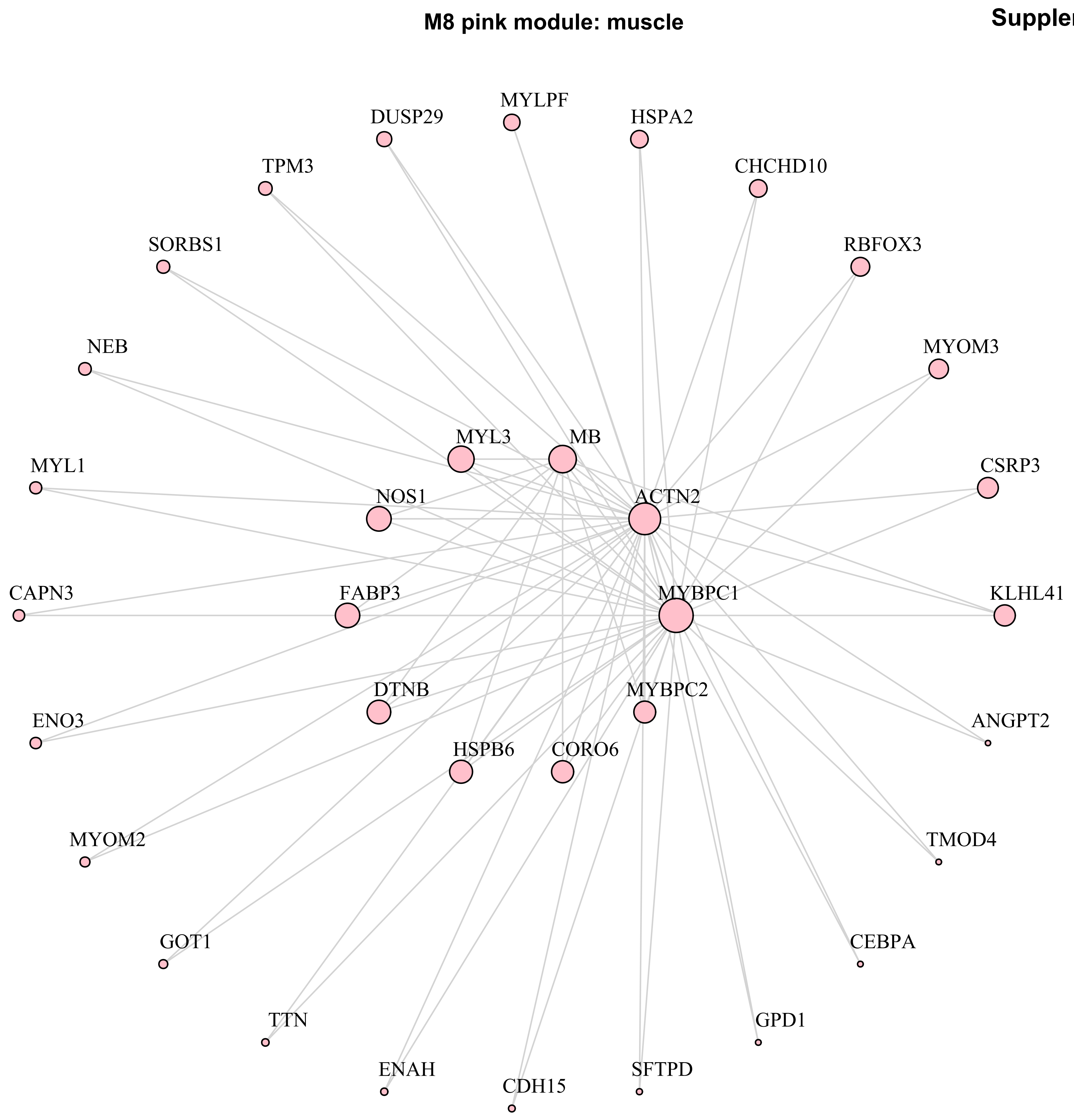

**Supplementary Figure 1**

**M4 yellow module: cell metabolism**

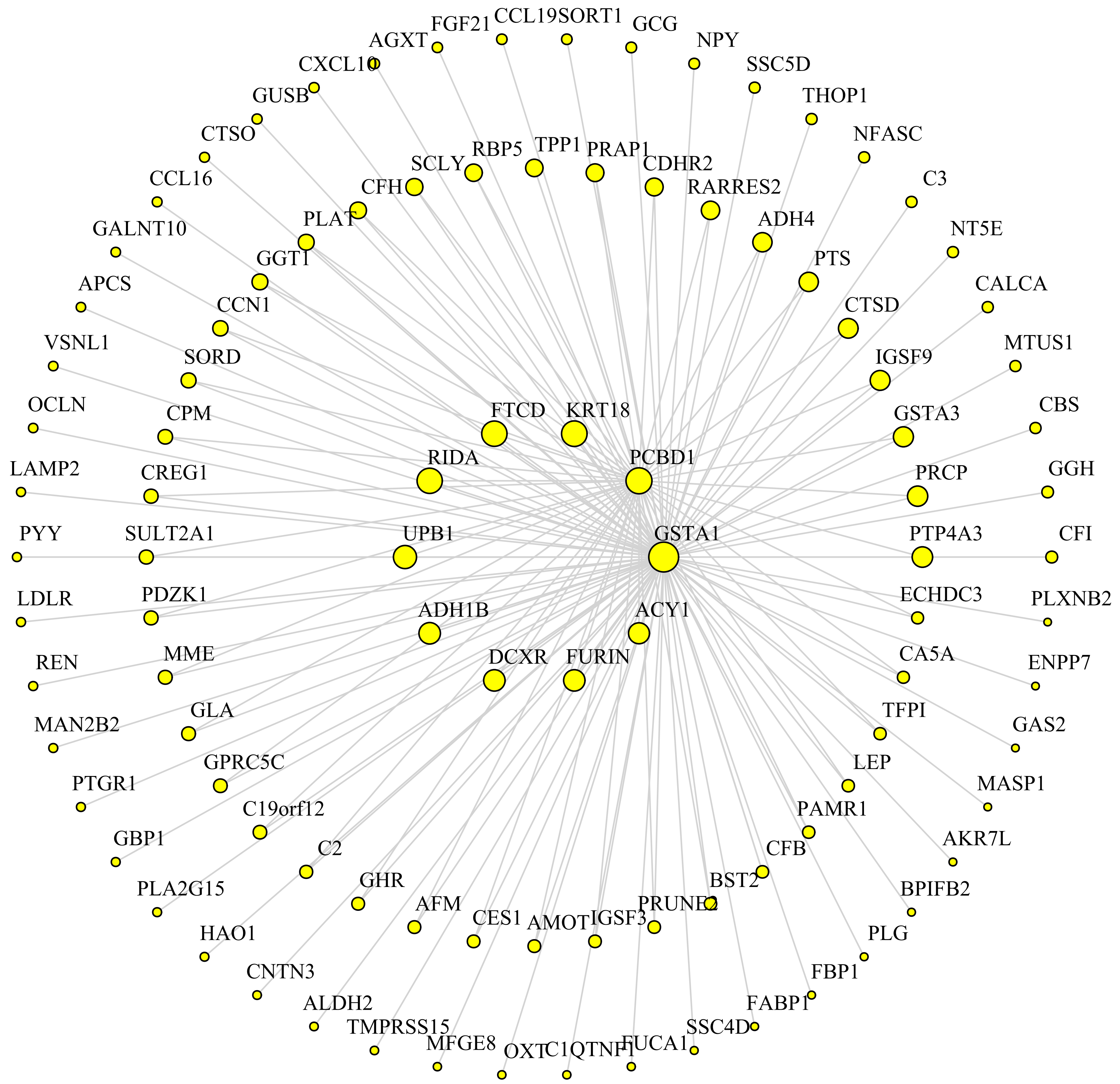

**Supplementary Figure 1**

**M2 blue module: TNF-signaling/cell adhesion**

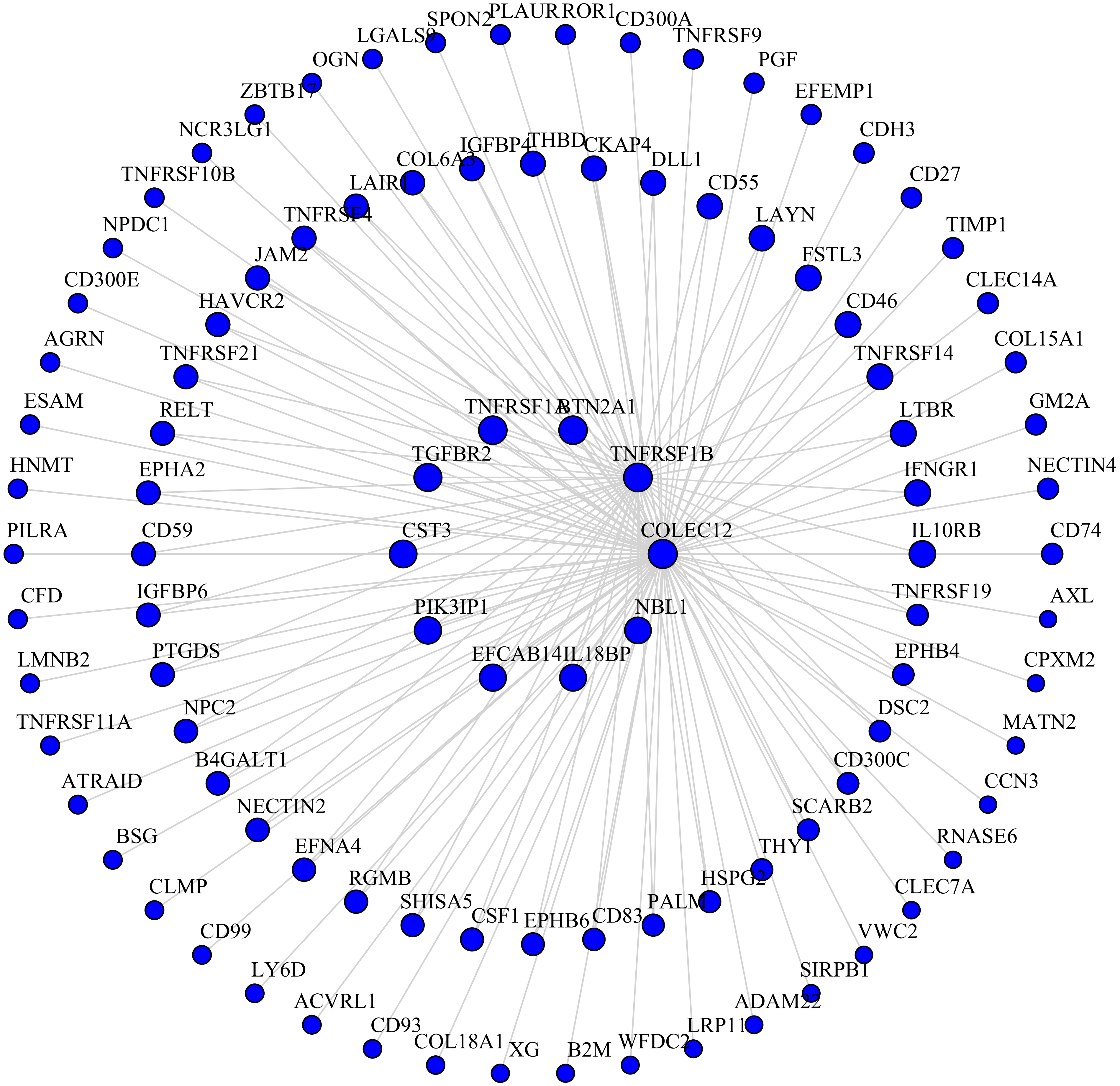

M5 green module: chemotaxis

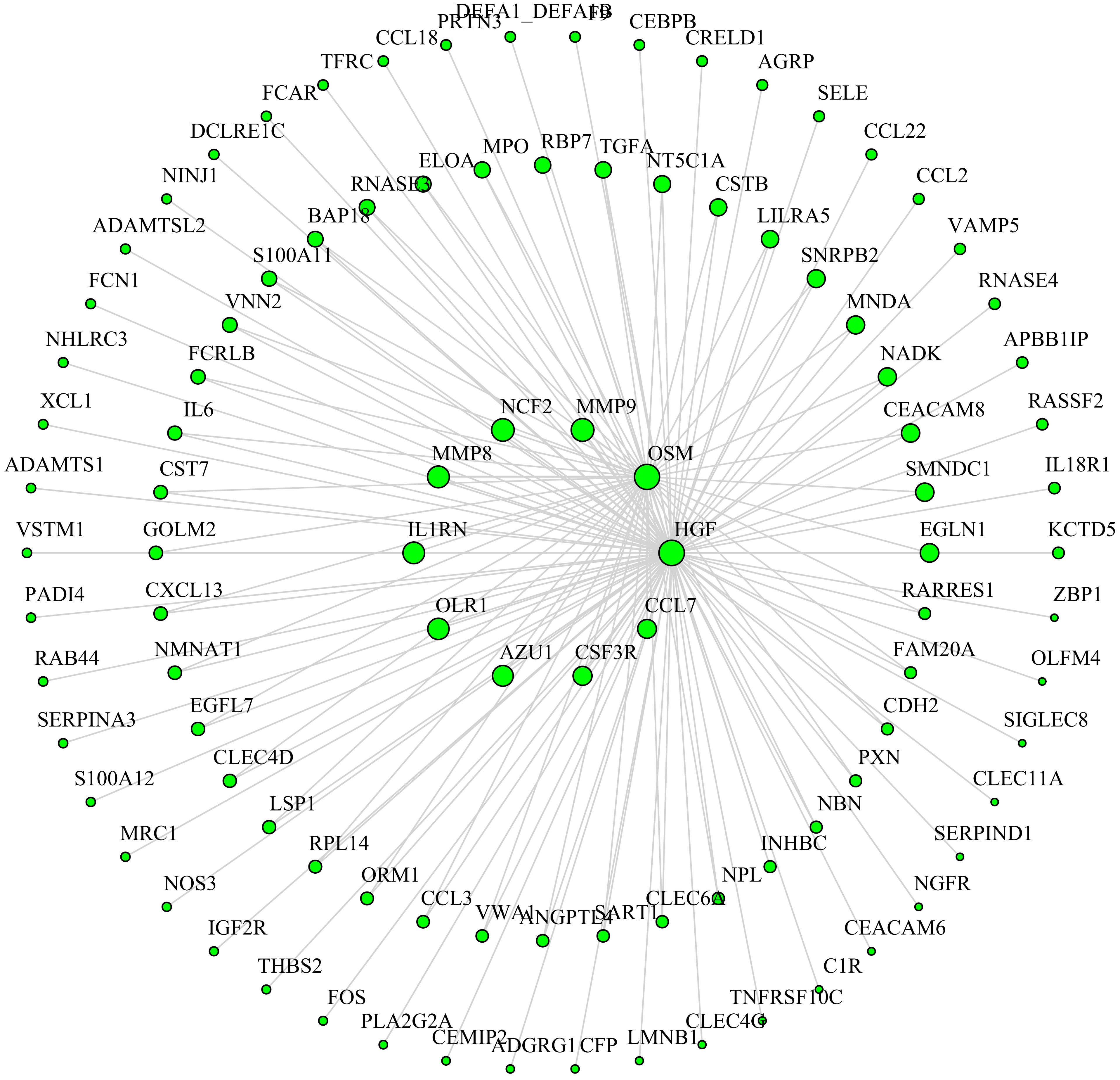

M6 red module: cellular detoxification

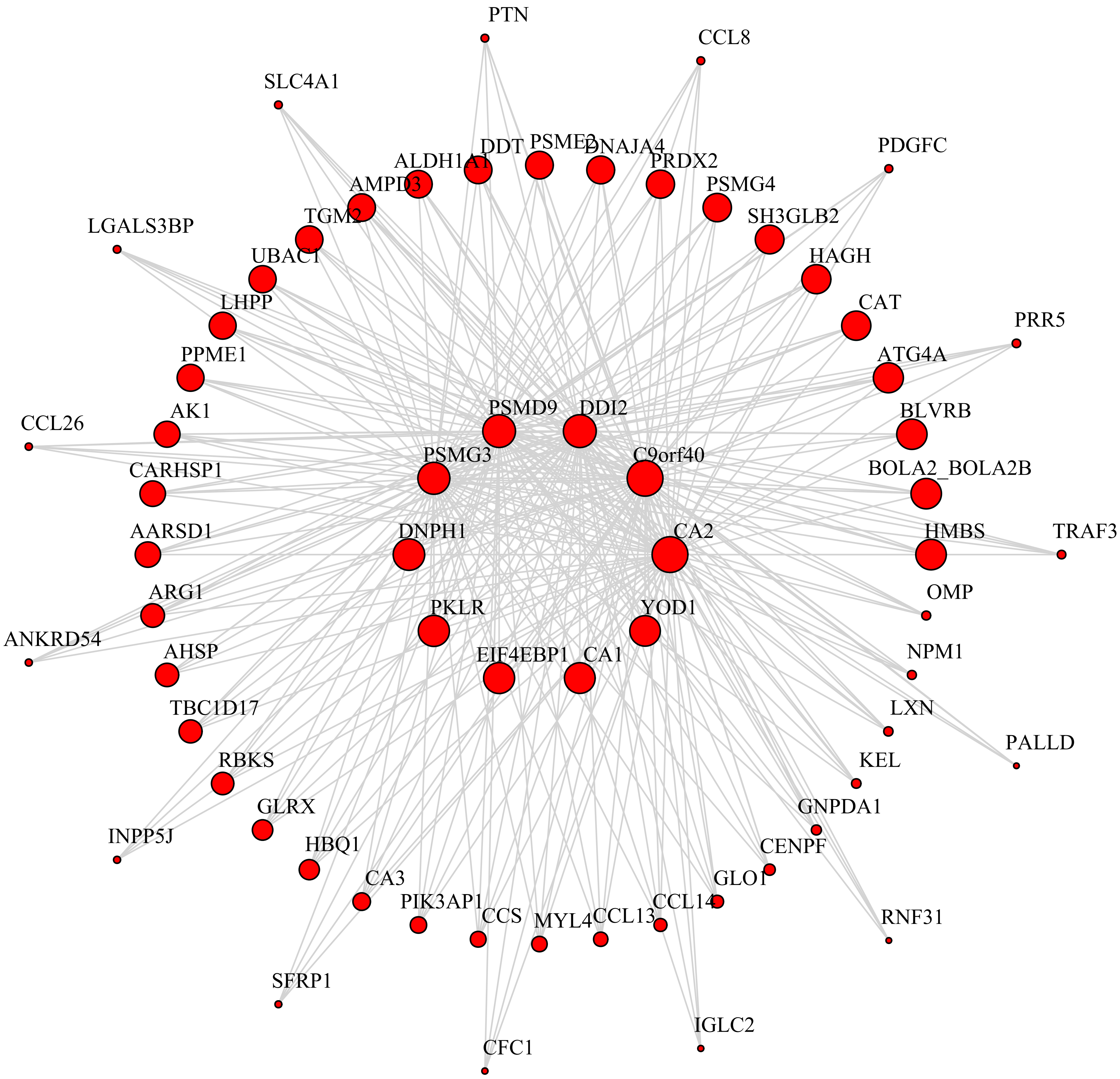

Supplementary Figure 1

**M1 turquoise module: intracellular signaling**

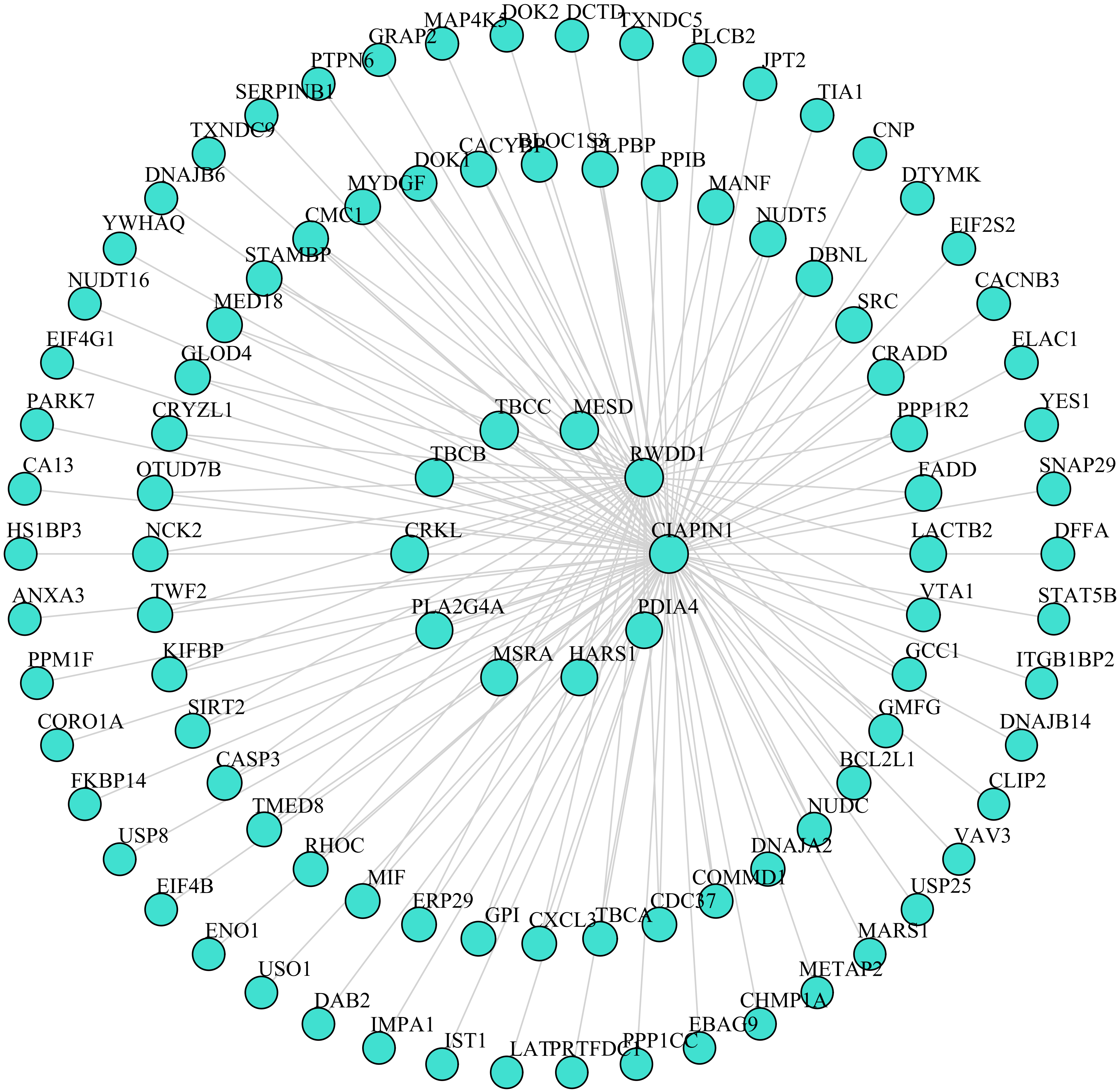
