## Supplementary figures and images for "Interrogating the plasma proteome of repetitive head impact exposure and chronic traumatic encephalopathy"

### Supplementary Figure 2

#### Supplementary Figure 2

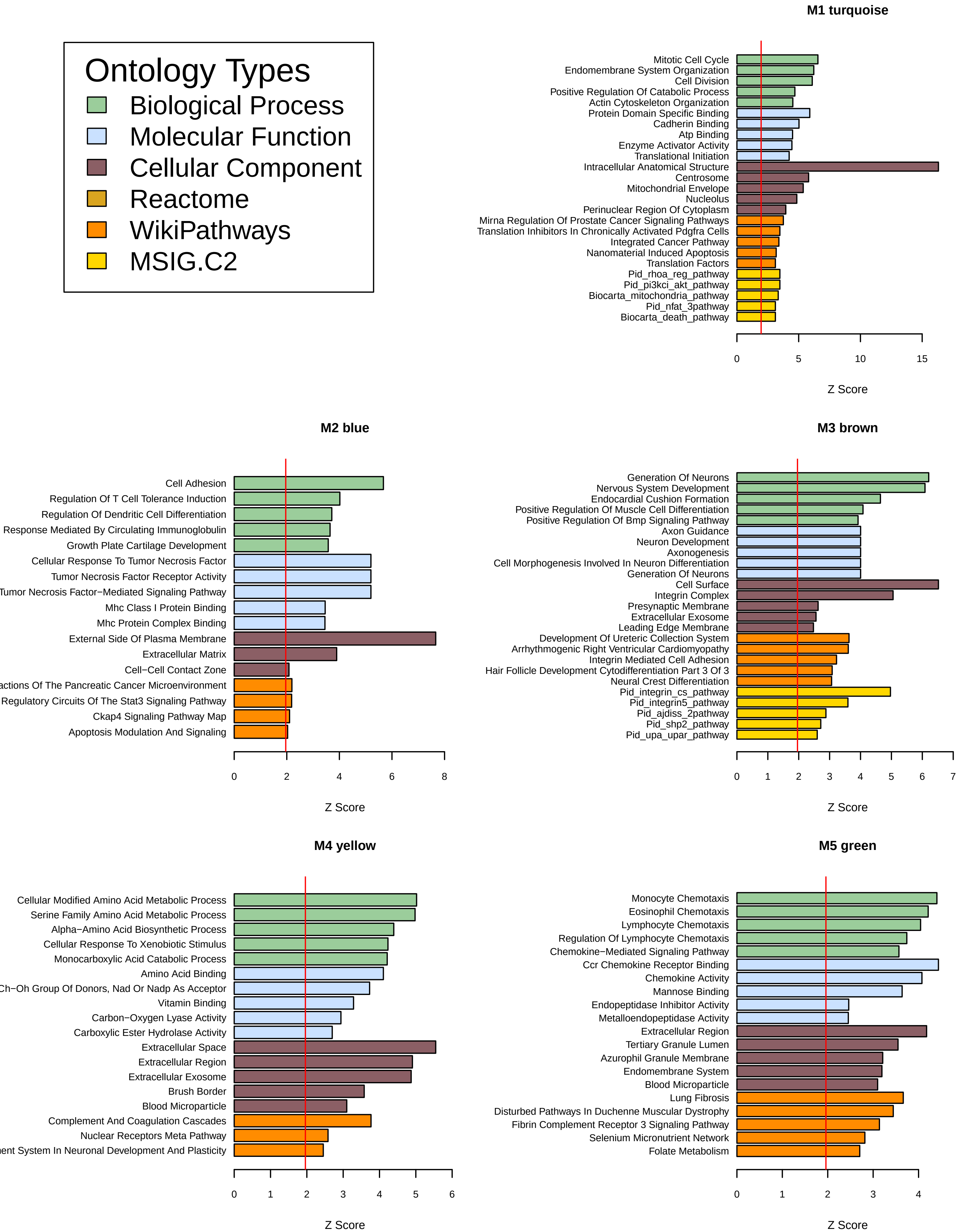

#### Supplementary Figure 2

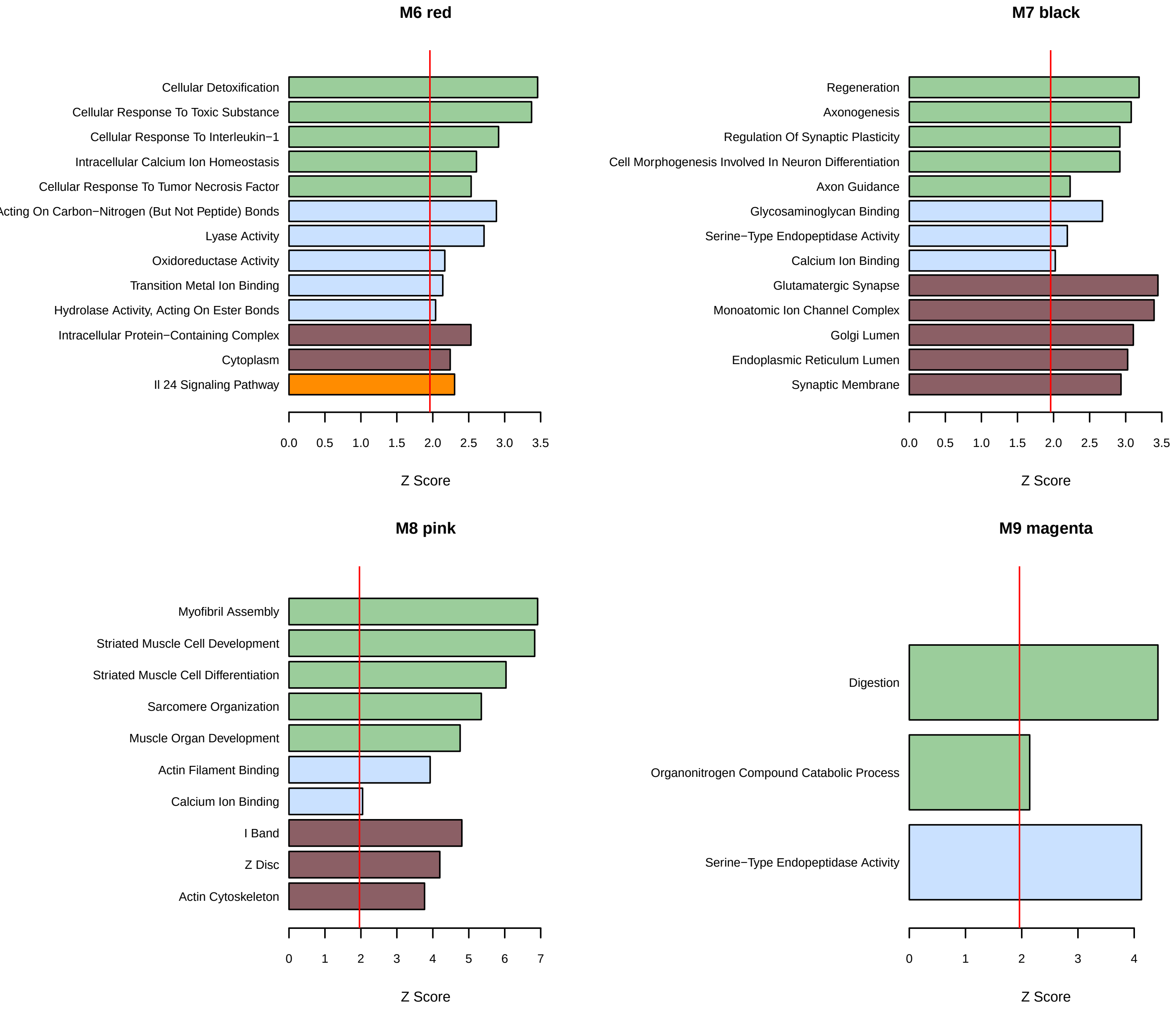
